# Sustained activation in basal ganglia and cerebellum after repetitive movement in cervical dystonia

**DOI:** 10.1101/2023.03.19.533030

**Authors:** John K. Kuster, Jacob M. Levenstein, Jeff L. Waugh, Trisha J. Multhaupt-Buell, Myung Joo Lee, Byoung Woo Kim, Guido Pagnacco, Stefano Sapienza, Paolo Bonato, Miriam L. Makhlouf, Lewis R. Sudarsky, Hans C. Breiter, Nutan Sharma, Anne J. Blood

## Abstract

**Introduction:** We previously observed abnormally sustained fMRI BOLD signal in the basal ganglia in focal hand dystonia (FHD) patients after they performed a repetitive finger tapping task. Focal hand dystonia is frequently a task-specific dystonia, where pathogenesis has been linked to excessive task repetition and aberrant sensorimotor plasticity. In the current study we asked if the effect of repetitive movement observed in FHD would be seen in other forms of dystonia where there is less known about the role of muscle use and repetition.

**Methods:** We evaluated fMRI BOLD signal time courses before, during, and after the block design finger tapping task in cervical dystonia (CD) patients for comparison with our previous study of focal hand dystonia. To ask whether brain function differences in CD might be linked to aberrant cervical muscle function during the task, we acquired surface EMG (sEMG) from a subset of the CD patients and controls at a follow-up session. Specifically, we measured hand/arm and cervical muscle activity during the tapping task.

**Results:** We observed abnormally sustained BOLD signal after tapping in CD participants in the left putamen and left cerebellum during non-dominant (left) hand tapping runs. BOLD signals in left putamen and cerebellum were also abnormally elevated in CD during the tapping task itself and escalated over time as the task was repeated. The laterality of sustained putamen signal was the same in CD as in our previously studied FHD cohort: patient/control differences were observed only in the left hemisphere. In contrast, there were no cerebellar BOLD time course abnormalities in the FHD cohort. sEMG signal across the left-hand tapping run bore strong resemblance to contralateral fMRI BOLD time courses at both the group and the individual level, including escalation of signal during tapping and sustained activity during rest blocks following tapping in CD. These effects were largest and most consistent in the right sternocleidomastoid (SCM) muscle. In contrast, task-related arm/hand muscle (flexor carpi radialis) sEMG was similar between CD and controls and did not escalate in either group.

**Discussion:** Based on these preliminary, qualitative findings, we propose that future paired imaging/EMG studies in larger cohorts are needed to evaluate the link between brain abnormalities and aberrant, task-related function in the dystonic muscles. While the imaging findings are not inconsistent with sustained brain activity reflecting an endophenotype and/or hemodynamic abnormality in CD, the sEMG data support the additional possibility that in CD there is an amplified response to the normal mechanical demand of hand tasks on proximal/core body musculature.

## Introduction

Dystonia is a neurological movement disorder characterized by sustained, involuntary muscle contractions. It may occur as a primary disorder or as a secondary symptom of other disorders such as Parkinson’s Disease. Traditionally, non-genetic, primary dystonias are classified by anatomical location of symptoms and by whether they are triggered by a specific task or not. Task-specific dystonias are thought to affect select motor programs such that the affected body region(s) can appear normal during rest and other tasks. It is believed that in many cases, repetitive, excessive use of the affected motor programs plays a role in the pathogenesis of task-specific dystonias such as writer’s cramp, musician’s dystonia, and laryngeal dystonia [1; 2; 3; 4; 5; 6].

In contrast to hand dystonias, such as writer’s cramp, cervical dystonia (CD) is not typically considered a task-specific dystonia. However, isolated reports of task-triggered CD do exist [7; 8; 9] suggesting there may be more to “use” in this form of dystonia than previously thought. Understanding the role of “use” or “overuse” of cervical muscles in CD would have substantial implications for improving prevention and treatment, including increasing our understanding of mechanisms behind sensory tricks and botulinum toxin treatment. A number of previous imaging studies have shown activation amplitude abnormalities in individuals with CD during hand or arm motor tasks, despite the fact that these tasks did not involve overt movement of the affected region. This alone does not address task-specificity but it does suggest that the brains of these individuals respond abnormally to “use” of the sensorimotor system, even when the dystonic muscles are not an obvious part of the task.

With the aim of studying the neural effects of muscle “use” in task-specific focal hand dystonia (FHD), we previously showed there was sustained elevation of blood-oxygenation-level-dependent (BOLD) activity in the basal ganglia after completing a bilateral repetitive finger tapping task [10]. These findings were not only consistent with pathophysiology triggered acutely by repetitive movement in these individuals but also suggested that reinforcement of motor programs can outlast performance itself, which could be one mechanism by which programs become “over-reinforced” to produce dystonia. The evaluation of temporal information in the previous study differed from other task-related imaging studies in this population because BOLD signal is typically collapsed over time and assumes that task performance does not influence subsequent blocks of imaging used as a control comparison (e.g., rest). With the aim of studying the effects muscle “use” in a focal dystonia that is not typically considered task-specific, the current study sought to evaluate whether the finger tapping task elicited sustained brain activation in CD as it did in FHD by evaluating whether the BOLD signal changed over the course of task execution itself.

In the context of the questions posed in the current study it is possible that some of the traditional segregation of dystonias into those that are task-specific versus those that are not task-specific reflects a semantic distinction, relating to differences in the way we use different anatomical regions and way we use the terms “task” and “use”. For example, task-specific dystonias such as spasmodic dysphonia and writer’s cramp involve anatomy that is used in an obvious and direct way to execute tasks (speaking, writing etc.). In contrast, muscles in the neck and trunk are not used in obvious ways to conduct “tasks” but they are nonetheless used almost constantly, often in the background [11; 12; 13; 14; 15; 16; 17; 18; 19; 20; 21; 22; 23; 24; 25; 26]. To some extent, then, the muscles in this region are “on task” much of the time.

The kinesiology literature predicts that movement itself is not necessary to recruit symptomatic muscle activity. For example, it is well established that when one part of the body initiates a movement (e.g., the upper extremities during manual tasks), there are adjustments in other parts of the body to maintain overall balance and stability (e.g., [14; 16; 17; 26; 27; 28]). This includes anticipatory postural adjustments but may also include other body stabilizing functions that help to implement motor control, which are not necessarily gravity/position dependent (e.g., axial function observed while performing a motor task in a supine position), and which vary in presence and amplitude even across healthy controls [26; 27; 28]. With the goal of intersecting the dystonia literature with kinesiology to interpret imaging findings, we asked in the current study whether the so-called “tasks” demanded of the cervical muscles tap into the affected motor “programs.”

While neural activity abnormalities outside the symptomatic body region in dystonia are consistent with endophenotypic traits (e.g., [29; 30]), subclinical pathophysiology (e.g., [30; 31]) and/or hemodynamic confounds to fMRI (e.g., [32]), in the current study we tested a complementary hypothesis that some of these effects may also reflect task-related dystonia symptom pathophysiology due to “beyond the arm/hand” mechanical demands of the finger tapping task. To do this we (1) evaluated activation magnitude and time courses in CD and controls in primary motor cortex regions outside the hand region and (2) brought back a subset of CD and control participants to evaluate surface electromyography (sEMG) of cervical muscles during the finger tapping task.

In the current study we also made two minor modifications and additions to our previous approach to answer further questions about our previous findings: First, to better understand the relationship of brain activity abnormalities to the task itself, the current study used a unilateral version of the tapping task to determine the laterality of movement that yielded these effects in dystonia and the laterality of the BOLD findings, e.g., were the findings always contralateral to the tapping hand? Second, we included longer rest blocks (60 seconds rather than the 30 seconds used previously) to increase the number of pre-tapping rest repetition time (TR) acquisitions and provide equal numbers of TRs during tapping and post-tapping blocks. We also added comparisons of cerebellar activation time courses to the current study, since this region has been increasingly implicated in the pathophysiology of dystonia [e.g., [33]] and may be useful in differentiating between different forms of dystonia [34; 35; 36; 37].

## Methods

### Ethics Statement

All participants included in this study signed written informed consent prior to participation in the study, and the study was approved by the Mass General Brigham Institutional Review Board (protocol #2006P000960). All experiments were conducted in accordance with the principles of the Declaration of Helsinki.

### Participants

#### Participant inclusion criteria

Twelve individuals diagnosed with primary cervical dystonia (CD) (4 males, 8 females, mean age = 54.58±8.68 years) and negative for the DYT1 mutation, were included in the current study. Three additional CD patients from an original pool of 15 were excluded based on criteria outlined below. CD participants were negative for all other neurological diagnoses (including essential tremor). Twelve healthy control participants, with no history of or current neurological or psychiatric disorders, and no treatment with neuroactive medications, were matched one-to-one to CD participants by age (within five years), gender, and handedness (see Table 1, mean age = 54.42±9.07 years). Controls were confirmed in the clinical screening (see below) to have no evidence of dystonia or other neurological or psychiatric disorders.

**Table 1:**
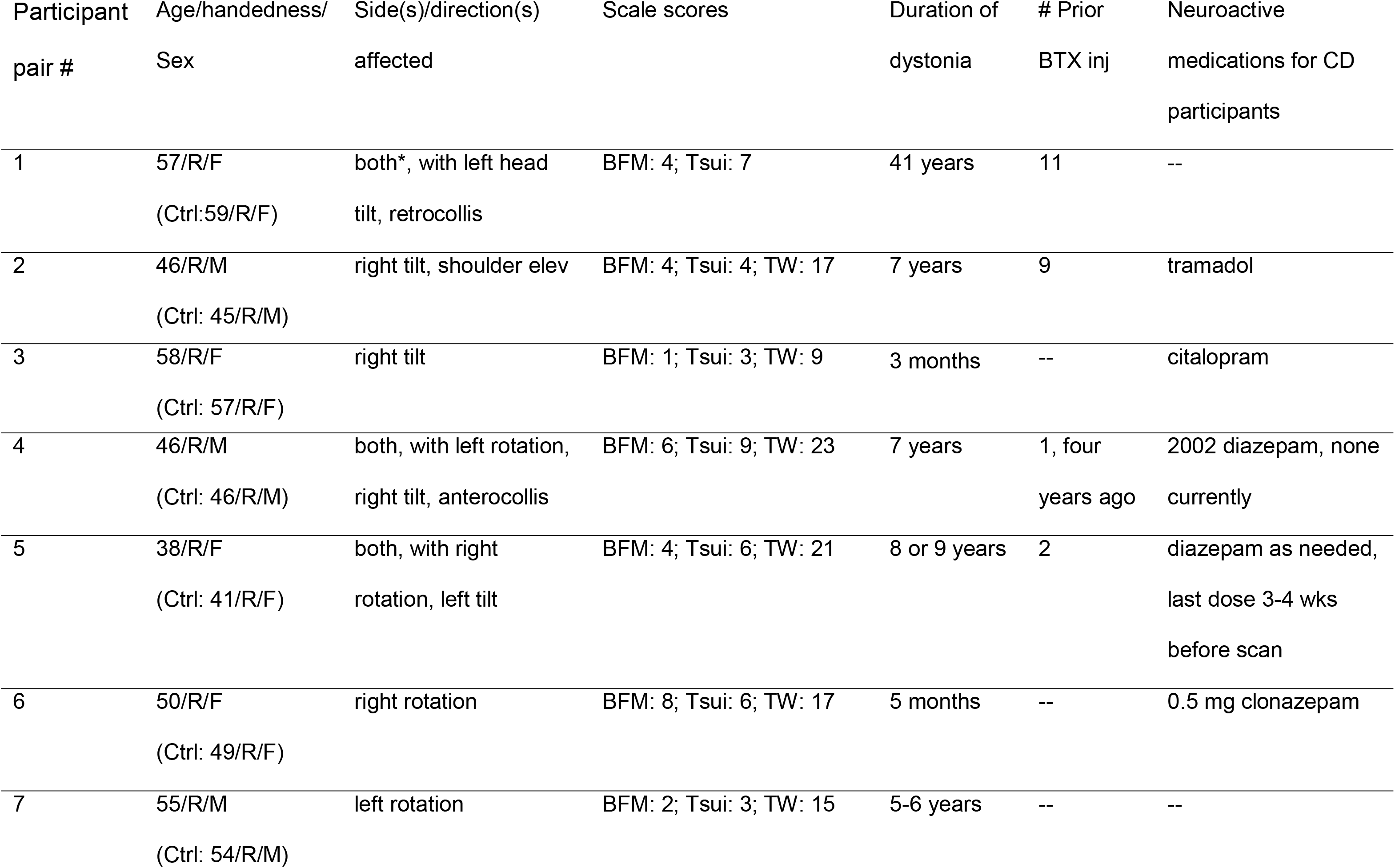
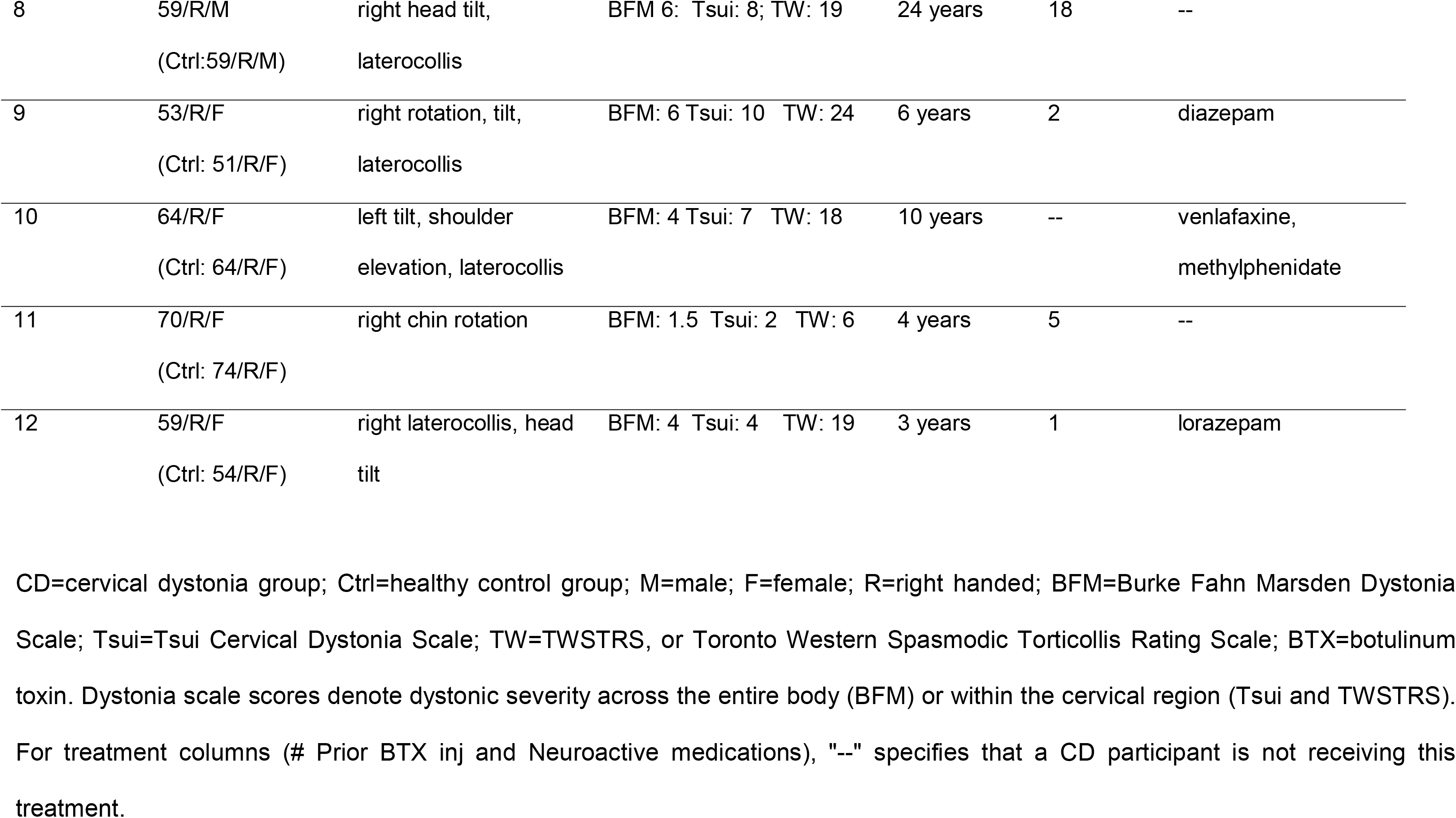
Demographics and clinical characteristics for CD participants and one-to-one matched controls.

As referenced in the Introduction, the model underlying our motor control research [34; 35] predicts that (1) pathophysiology differs across at least four qualitatively different types of dystonia symptoms [34] and (2) the fMRI abnormality we wished to detect would not be expected to be a component of pathophysiology in (and therefore not observed in) individuals exhibiting purely "static" dystonia symptoms at rest, we excluded from an original pool of 15 CD patients and 15 matched controls any potential CD participants who exhibited ‘static” symptoms at rest. We identified “static” only patients as those meeting one of the two following criteria on the Toronto Western Spasmodic Torticollis Rating Scale (TWSTRS) CD rating scale [38; 39]: (1) a score of 2 or higher for [limited] range of movement or (2) a score of 1 for range of movement AND little to no effect of sensory trick [i.e., a patient-performed action or sensory stimulus that reduces dystonia symptoms [40]]. While these rating scale criteria do not perfectly distinguish different types of dystonia symptoms, they are the best validated and objective measures currently available. Further discussion of differentiating factors of "dynamic" and "static" dystonia symptoms can be found in [41]; note that in the Muller publication the terms, "dynamic" and "static" are referred to as “phasic” and “tonic”, respectively. Based on these criteria we excluded three patients from our original pool of 15, as well as their matched controls, resulting in twelve CD patients and twelve matched controls for the analyses reported here. Note that most individuals with dystonia have a mixture of dynamic and static symptoms and the presence of static symptoms would not be expected to interfere with the effect we were aiming to detect— we aimed for the presence of dynamic symptoms, rather than the exclusion of static symptoms to test our hypotheses.

The scan session was timed such that any patients receiving botulinum toxin injections to treat dystonic symptoms were scanned at the end of a treatment period, at least three months after injections and within one week of their next set of injections. This timing was used for both the imaging session and for follow-up sEMG sessions.

#### Clinical screening

Immediately prior to the MRI session, all participants completed a clinical history (including past and current medications) and a clinical assessment, including standard neurologic exam and completion of standardized dystonia severity scales (Burke Fahn Marsden [BFM; [42]], Tsui [43], and Toronto Western Spasmodic Torticollis Rating Scale [TWSTRS; [38; 39]]) by a movement disorders physician (N. Sharma). Table 1 characterizes the demographics and clinical profiles for each individual with CD, including duration and severity of the disorder and past or current medication status. For those individuals who were currently being treated with botulinum toxin (BTX) injections, scanning was conducted at the end of a treatment period (within the week before the next scheduled injections) so that clinical benefits of botulinum toxin were at a minimum and patients’ dystonia presentation reflected their least treated state.

### Data acquisition procedures

#### Motor task and fMRI block design

For this fMRI study, all participants performed a modified version of the finger tapping task published previously [10]. As before, participants alternated tapping the index and little finger (digits 2 and 5) on button boxes equipped with four tandem, computer-sized keys. Tapping was aurally cued at a 2Hz frequency for three, 60-second blocks, with 60-second rest blocks preceding and following each tapping block (block design shown in green in Figure 1); task cue presentation was conducted using Presentation software (Neurobehavioral Systems Inc., Berkeley, CA). TR length remained at 2.5 seconds as in our original study [10]; however, the duration of all four rest blocks was expanded from 30 seconds in our previous study to 60 seconds in the current study. This adjustment increased the number of pre-tapping rest acquisitions and resulted in an equal number of acquisitions during tapping and post-tapping blocks. Participants were instructed to relax and not move their hands during rest blocks. Keypresses and hand movement were recorded throughout each run. Padding was placed beneath each participant’s upper arms and elbows so that he/she could reach the button keys with arms supported, thereby minimizing task-related head movement and neck strain. Participants were also instructed to close their eyes during the entirety of each run.

**Figure 1.**
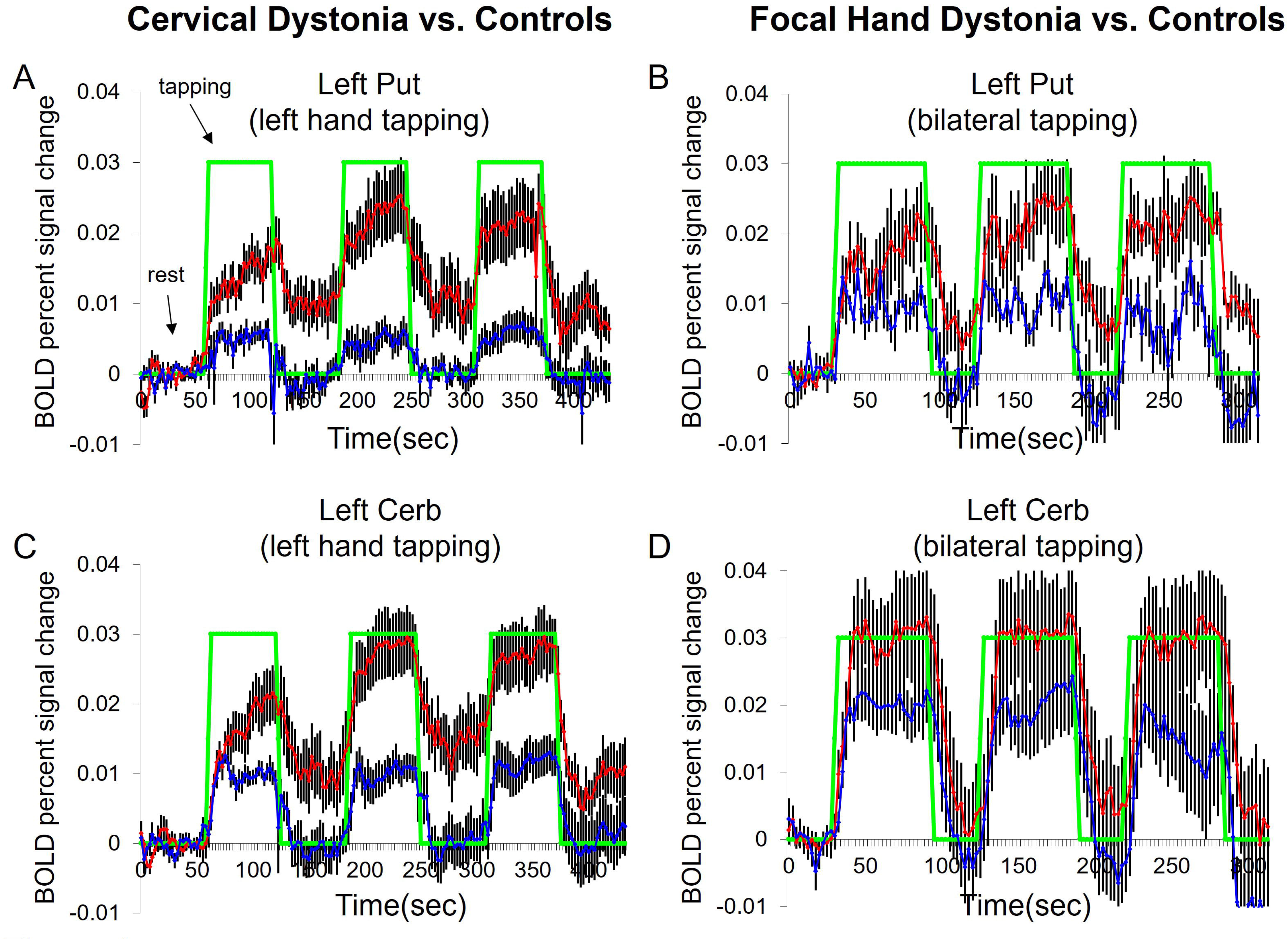
**(A)** BOLD time course comparisons in left putamen, cerebellum, and primary motor cortex dorsal to the hand representation for left-hand tapping. Red points indicate CD time courses and blue points indicate control time courses. Data are expressed as percent signal change (in decimal form) from the initial, pre-tapping rest block over time. The block design for the task is denoted by the green line on the putamen time course; total run time was 435 seconds, or seven minutes, 15 seconds (with 60 second rest blocks). Error bars represent standard error of the mean. **(B)** BOLD time course comparisons in right putamen, cerebellum, and primary motor cortex ventral to the hand representation for left-hand tapping. Red points indicate CD time courses and blue points indicate control time courses. Data are expressed as percent signal change (in decimal form) from the initial, pre-tapping rest block over time. The block design for the task is denoted by the green line on the putamen time course; total run time was 435 seconds, or seven minutes, 15 seconds (with 60 second rest blocks). Error bars represent standard error of the mean.

In addition to longer rest block time, the task was modified in the following ways from the earlier study [10]: (1) Participants performed unilateral (i.e., one hand at a time), rather than bilateral finger tapping, to evaluate findings in relation to laterality of movement. Each run involved tapping using only one hand for all three tapping blocks in the run; two runs were acquired for each hand, for a total of four task fMRI runs. Order was pseudorandomized across subjects to control for repetition effects. Because control participants were matched one-to-one with patients, we matched run order for each patient/control pair. (2) Participants did not make judgments of acute dystonia severity in the current study, to rule out that elevated signal in the previous study might have related to potential cognitive/evaluative processes involved in making such judgments during scanning. Furthermore, we evaluated a broader set of sensorimotor regions than in the previous study and evaluated one additional region from the original FHD cohort (the cerebellum) for comparison with findings in the CD cohort.

#### EMG task design

Surface EMG was acquired offline in six of the fMRI cohort CD patients and five of the healthy controls using the same button box finger tapping task design described above for fMRI. Participants lay in a supine position, identical to the position used in the MRI magnet. Run order for task laterality (left-versus right-hand) was identical to the order used for fMRI for each participant; for example, if the order was left, right, right, left for fMRI acquisition in a given individual, that same order was used for offline EMG runs. Thus, any order effects in the scanner within-subject would also be present during EMG runs and run order was pseudorandomized and counterbalanced across participants.

#### Button box data (finger tapping behavior measures)

During fMRI, any keypresses made were recorded and saved with a timestamp in Presentation software (Neurobehavioral Systems Inc., Berkeley, CA). These data were used to evaluate if participants performed the task as instructed (i.e., that they tapped during tapping blocks and did not tap during rest blocks) and to determine if there were task performance differences between CD and controls.

#### Hand motion sensors

Piezoelectric accelerometers (referred to as "motion sensors") were used to monitor hand movement to verify that participants were not moving their hands during rest blocks [10; 44]. Measures were acquired and recorded using LabView software (National Instruments, Woburn, MA). Accelerometers were Murata PKM11-4A0 (Murata Electronics North America, Smyrna, GA) with a silicon filling to reduce sensitivity to acoustic noise. Accelerometer data were acquired at a resolution of 200 acquisitions per second, using LabVIEW software and a National Instrument Data Acquisition Card (Model 6024E) on a Sony Vaio laptop.

#### Magnetic Resonance Imaging Methods

A 3.0 Tesla Siemens Tim Trio magnet system (Siemens AG, Medical Solutions, Erlangen, Germany) was used to acquire whole-brain echo planar imaging (epi) sequences (fMRI) while participants performed the finger-tapping task. Images were acquired using 5mm slice thickness with no gap and interleaved acquisition; this corresponded to 30 slices to cover the entire brain. Within plane (x/y dimensions) resolution was 3.125 mm x 3.125 mm. The repetition time (TR) was 2.5 seconds, resulting in 24 TRs/full volume acquisitions across each 60-second block. The echo time (TE) was 30 ms. High-resolution T1-weighted anatomical scans were acquired for registration purposes and anatomically-guided fMRI time course extraction. During all image acquisition, participants’ heads were stabilized as described previously [e.g., [10]] with foam padding around the head, and paddles on either side. All CD participants were able to lie still without experiencing head movements from their dystonia since lying supine substantially reduces symptoms in many patients. Any patients with remaining symptoms were allowed to position their head slightly off center if this enabled them to keep their head still throughout the scanning session. We verified the absence of head movements in patients using head movement analyses described in Supplemental Methods. Both T1 and epi (fMRI) sequences were acquired using Autoalign software [45] such that slice prescriptions were anatomically consistent across participants.

#### EMG acquisition parameters

Surface electromyography (sEMG) was acquired on a separate day from imaging and was performed in one of the clinical research exam rooms. Participants were positioned identically to the position in the scanner—on a padded exam table, in a supine position—and used the same button boxes and auditory cues to perform the same tapping task. During each run, sEMG was acquired using a Delsys 16-channel intraoperative EMG system. Data were acquired at a sampling rate of 8,000 Hz, a hardware gain of 10,000, a signal amplification, and passband gain of 5,000 for analog-to-digital conversion. Data were acquired using a paired electrode, differential approach for each location of interest. sEMG was preferred to needle EMG for the purposes of this study, to capture a snapshot of overall cervical load from the task, rather than measuring features at a single focal point.

Electrode Placement: A diagram of electrode placement is included as supplementary Figure 3. We acquired sEMG from the flexor carpi radialis to capture activity of hand/finger flexors. We acquired sEMG from the following cervical muscles, including muscles frequently injected with botulinum toxin in CD: sternocleidomastoid (SCM), splenius capitus, trapezius, and levator. For most channels, the first electrode of the differential was placed in the middle of the muscle (for flexor carpi radialis, SCM, and splenius capitus muscles). A single channel captured trapezius and levator muscle function together by placing the electrode just above the point of insertion of the levator at the scapula. The rationale for the placement and pairing of these two muscles is that the levator is closer to the surface at this point; if measured in the middle of the levator muscle (side of the neck) signal would be difficult to detect. In all cases the second (differential) electrode was placed one inch below the first electrode. Electrode location/placement was aided by asking subjects to briefly perform movement that involved the muscle(s) of interest.

### Data Analyses

#### 1. Tapping task performance during fMRI acquisition

##### Button box tapping data

We evaluated button box tapping data: (1) to confirm that all participants performed the task correctly and verify that they did not tap during rest blocks and (2) to compare patient and control tapping performance during tapping blocks. For analysis of tapping performance during tapping blocks we compared total number of taps (button box keypresses) across CD and control groups to verify that all participants performed the task properly, and that overall task performance (i.e., number of keypresses) did not differ across groups. Comparisons were made across groups for the total number of keypresses across all three tapping blocks (two-tailed t-test), as well as by block number (a 2 x 2 repeated measures ANOVA evaluating the interaction between group and tapping block number). Results are reported as uncorrected p values. For these negative control analyses, we required that the uncorrected p was at least 0.1 or greater to denote a negative result (i.e., no differences between groups).

##### Hand motion sensor data

We evaluated piezioelectric accelerometer (hand motion sensor) data for CD participants and controls to show that post-tapping rest block signal did not differ from pre-tapping rest block signal in CD or control groups. This negative control step verified that fMRI findings did not simply reflect differences in the relative amount of pre-versus post-tapping rest block hand movement in one or both groups.

Motion sensor preprocessing: Raw motion sensor recordings were acquired throughout each fMRI run. Raw data were both frequency and amplitude filtered to remove regular frequency artifacts produced by the magnet field. We high pass filtered at 0.5hz and low pass filtered at 5hz. After filtering, we normalized motion sensor data from each run to the rest block portions of the dataset for comparisons across participants and groups. The motion sensor data was similar to BOLD signal in that it measured relative, rather than absolute, signal amplitude across runs and sessions, and therefore, the relevant information for comparisons across groups came from relative signal during tapping versus rest blocks. We normalized to signal amplitude (peak-to-trough) across all four rest blocks for a given run (see Supplementary Methods). After normalization, runs were averaged for a given participant/hand/condition before making group comparisons.

Motion sensor statistical comparisons: Once motion sensor data were normalized, we verified there were no CD/control differences in hand movement in either hand during pre-tapping versus post-tapping rest blocks, for any condition that showed CD/control differences in BOLD signal. Specifically, we assessed whether there were any group (CD versus control) by condition (pre-versus post-tapping rest) interactions using a 2 x 2 repeated measures ANOVA, with condition (pre-versus post-tapping rest) as the repeated measures factor, and average BOLD signal for each participant and event. We also ran post hoc paired samples t-tests comparing pre-tapping rest to post-tapping rest separately within each group and for each hand, as described in the fMRI group *Contrast Analyses* section below, for the condition showing fMRI group differences. Thus, comparisons were analogous to the comparisons made for BOLD signal.

##### Head movement during imaging

In addition to standard procedures for head stabilization during data acquisition and motion correction during fMRI analyses, we used two further approaches to verify that head movement was not a factor in our findings. Please see Supplementary Methods for descriptions of these analyses and their results.

#### 2. fMRI analyses

fMRI and structural MRI data were analyzed using FreeSurfer analysis software (http://surfer.nmr.mgh.harvard.edu), including contrasts, segmentation, and time course extraction. Note that in our previous study [10] we computed brain activation by summing BOLD signal across regions of interest instead of using a standard voxelwise analysis to address previous evidence that activation is often broader or less consistently localized in patient populations (including dystonia) than in healthy controls [46; 47; 48; 49; 50; 51; 52]. This factor can lead to false negatives when calculating group differences using a standard voxelwise analysis. In addition, it is important to note that typical analyses would model out some or all of the effects we observed, assuming they were non-physiological signal drift. The amount of across-run drift correction differs across fMRI analysis platforms and some model out all drift while others model out only parts of it. Therefore, we used the same analysis platform (Freesurfer/FSFAST) used in our previous study to allow direct comparison with our previous findings in hand dystonia patients.

Group statistical comparisons were identical to those used in our previous publication in a FHD population [10], except that we evaluated additional regions in the sensorimotor network in the current study. In addition, group-level contrast analyses were done using 2 x 2 repeated measures ANOVAs and within-group post-hoc analyses were done using paired samples t-tests, rather than the Mann-Whitney test used in our previous publication, due to our larger sample size.

##### Image preprocessing

fMRI images were preprocessed using standard procedures in the Martinos Center FS-FAST fMRI analysis stream (http://surfer.nmr.mgh.harvard.edu/fswiki/FsFast). This included motion correction, intensity normalization, and spatial smoothing. These steps were conducted as described previously [10].

##### Individual contrast analyses and fMRI time course extraction

Contrast analyses and time course extraction were conducted using procedures described previously [10], including standard Freesurfer FS-FAST processing of whole brain fMRI BOLD signal within each participant in native space, to identify voxels associated with tapping versus rest, using a general linear model (GLM) and standard t-test analysis (http://surfer.nmr.mgh.harvard.edu/fswiki/FsFast). Also as described in our previous study, after these contrast analyses were conducted, masks for our regions of interest (ROIs; see next paragraph) were overlaid with contrast maps on preprocessed fMRI maps (still in native space) to extract average fMRI signal time course across voxels meeting a certain activation threshold within each ROI (t=6.0 for cortical ROIs; t=3.5 for subcortical ROIs). As described previously [10], we used a higher threshold for cortical ROI time course extraction because cortical sensorimotor regions activate so robustly to movement that we predicted group amplitude differences would be best detected by extracting signal at the higher end of the signal range. For cortical ROIs hypothesized to encode activity relating to cervical muscle function, we used the subcortical threshold of t=3.5 because we expected these regions to activate less robustly.

*ROIs evaluated:* In the current study, we extracted BOLD time courses from 15 brain regions (ROIs), separately for left and right hemispheres and for left-and right-hand tapping, with two regions hypothesized, *a priori*, to show group differences. These *a priori* ROIs were (1) putamen and (2) cerebellar cortex. The other ROIs were components of sensorimotor circuitry commonly evaluated in movement disorder studies [e.g., [53; 54; 55; 56; 57; 58; 59], including (3) globus pallidus, (4) caudate, (5) thalamus, (6) primary motor cortex overlying hand representation (1°Mh), (7) areas 3a and (8) 3b of primary somatosensory cortex overlying hand representation, (9) supplementary motor area, (10) dorsal premotor cortex, (11) ventral premotor cortex, (12) area 1 of somatosensory cortex, and (13) area 2 of somatosensory cortex. In addition, for post hoc analyses of motor cortex neck representations we evaluated primary motor cortex (14) dorsal (1°Md), and (15) ventral (1°Mv) to the hand representation [60] for evidence of two neck representations in motor cortex, including the less frequently discussed dorsal region). Cortical ROIs were defined as described in Blood et al., 2004, based on sensorimotor segmentation of inflated brains [61]. Subcortical ROIs were defined by automated segmentations generated by the Freesurfer “recon-all” command, using Freesurfer version 5.3. Please see Supplementary Methods for description of 1°Md and 1°Mv segmentations, which excluded the hand representation). Please see Supplementary Figure 1 and main text Figures 3 and 4 for images of the ROIs.

##### Preparation of fMRI time course data for visualization and group analyses

Time course data were prepared for visualization and statistical contrast analyses using the following procedures. For the purposes of visualizing time courses across each group in our figures, percent signal change was calculated for each TR of the time course relative to average signal from the first (pre-tapping) rest block. For contrast analyses described below (repeated measures ANOVAs and paired-samples t-tests), BOLD values for each TR were normalized to the average BOLD value across the entire run, rather than to the pre-tapping rest block, to avoid introducing artificial signal homogeneity across participants in the pre-tapping rest block. As in our previous study [10], for all rest block contrast analyses, TRs during the signal decay period (first six TRs) of each rest block following tapping were excluded to avoid contamination of the rest block signal with task-related BOLD response. The six transition TRs between tapping and rest blocks were also excluded. To align statistical power (i.e., number of TRs) across all rest blocks, we also eliminated the first six TRs from the pre-tapping rest block. The remaining datapoints were averaged across block(s) for each condition (i.e., pre-tapping rest, tapping, and post-tapping rest) for entry into group contrast analyses.

*Group contrast analyses (CD versus control) of fMRI time course data to test for sustained signal in CD following tapping (**pre-tapping rest versus post-tapping rest**), and elevated signal during tapping itself (**pre-tapping rest versus tapping blocks**)*

Contrast analyses were conducted in the following way: (1) For each ROI being evaluated and for each of the two hands (left-hand and right-hand) we tested whether there was a group (CD versus control) by condition effect for the following two conditions: (a) post-tapping rest BOLD signal relative to pre-tapping rest signal and (b) tapping signal relative to pre-tapping rest signal. This was done using 2 x 2 mixed models ANOVAs, with group as the between-subjects factor and condition as the within-subjects factor. *A priori* ROIs (putamen and cerebellum) were run in a first tier of analyses before running analyses for the remaining ROIs. (2) For any significant group by condition effects in the mixed models ANOVA, we then ran post hoc paired samples t-tests (two-tailed) for the *a priori* ROIs (putamen and cerebellum), comparing pre-tapping rest to post-tapping rest in the CD group. For all contrasts, left and right hemisphere data were analyzed separately.

##### Group contrast corrections for multiple comparisons

Corrections for multiple comparisons were made in the following way. For *a priori* ROIs, the first tier of analyses involved a total of eight comparisons (2 ROIs x 2 task laterality (left/right hand) x 2 hemispheres), so we used a Bonferroni correction of p=0.05/8 = 0.00625 as our significance threshold for ANOVAs in *a priori* ROIs. Since only one of the two hands (left-hand) showed CD/control differences, we ran the pre-versus post-tapping post hoc t-test comparisons in the putamen and cerebellum for the left-hand runs only. These post hoc tests were run only for CD, with a total of four comparisons (1 group x 2 ROIs x 1 task laterality x 2 hemispheres), and a p=0.05/4 = 0.0125 threshold for correction.

For the remaining, non-*a priori* sensorimotor regions, there were a total of 13 ROIs for which we ran ANOVAs for each hand and each hemisphere separately, leading to a corrected p value of p=0.05/(13 x 2 x 2) = 0.000962. This correction was used for post-tapping rest comparisons and for tapping comparisons.

For all comparisons described above, contrasts yielding p values within an order of magnitude of the corrected p value were considered trends. That is, p≤0.0625 for *a priori* ANOVAs, p≤0.125 for paired t tests, and p≤0.00962 for all other brain regions,

##### Analysis of signal “escalation” in fMRI time courses during tapping (quantitative testing of qualitative observations made in Blood et al., 2004 [10])

Because we previously [10] made a qualitative observation that there was progressive signal increase over the course of individual tapping blocks in the ROIs that also showed sustained post-tapping function [10], we tested this *a priori* hypothesis quantitatively in our CD cohort. We compared BOLD signal at the beginning versus end of each tapping block to determine if there was a difference in CD but not in controls. We excluded the first three TRs of each tapping block for this analysis to remove normal signal rise time at task onset; this 7.5 second time period (3 TRs x 2.5 seconds per TR) fully covered signal rise time for both groups. We then compared the average (across TRs, then across tapping blocks, then across runs for each participant) BOLD signal for the fourth through seventh TRs of each tapping block (“beginning” TRs) to the average BOLD signal for the last four TRs of each tapping block (“end” TRs). We then ran a two-tailed, paired t-test to evaluate the hypothesis that there would be greater signal at the end than at the beginning of each block in CD, but not in controls. We ran this test only for the two *a priori* ROIs that showed significant group differences during tapping in our initial contrast analyses (left putamen and left cerebellum). With four comparisons (two groups and two brain regions per group) we required a Bonferroni correction for multiple comparisons of p=0.05/4 = 0.0125 to reach statistical significance.

##### fMRI time course analysis in non-activated regions to rule out scanner signal drift (negative control analysis)

All fMRI analysis platforms use algorithms to remove effects of potential scanner drift, but each platform uses slightly different algorithms. The FS-FAST platform used the algorithm described by Worsley and colleagues [62] to remove drift from the BOLD signal output in the current and previous study. Because BOLD signal drift cannot be easily distinguished from biological signal changes over time by drift algorithms we conducted the analysis used in our previous study [10] to gain further confidence that our findings were not due to residual magnet or head movement-related drift, using the following procedure to show that there was no global drift across the brain. For ROIs showing significant group differences (i.e., group x task/condition interactions), BOLD signal time courses were extracted for voxels that were non-significant in tap versus rest contrast results (i.e., -1.5<t<1.5), and ANOVA and t-test analyses were run as described above for activated voxels, comparing group x condition (pre-versus post-tapping rest) interactions, and within-group evaluations for differences in pre-versus post-tapping rest BOLD signal. For these negative control analyses we required that p > 0.1.

##### Post hoc questions: Evaluation of motor cortex outside the hand region during tapping in both groups (1°Md and 1°Mv)

One of our post hoc questions was whether there was indirect evidence that cervical muscles were used during performance of the finger tapping task. We asked this question by evaluating regions of motor cortex where neck representation might be expected. This was not a group contrast question but, rather, a question whether the task recruited cervical muscle function in either group. Because our initial analyses were designed to extract BOLD signal for amplitude comparisons between patients and controls, the methods for extracting time course data used *t* thresholds for data within-subjects, rather than *p* thresholds determined to be significant activation across a group. Therefore, it was not legitimate to use the time course data to determine whether there was group “activation” in a given region using this approach (as opposed to group differences in amplitude) as this would have been circular. For this reason, the specific question about neck representation function (location and time course features) was designed as a qualitative evaluation only; we propose the need for future studies designed to test, *a priori*, whether there is significant activation in these regions.

##### Post hoc questions: fMRI analyses of the cerebellum in our previous FHD cohort

After the CD cohort analyses were conducted, we revisited our FHD data to evaluate whether the effects we observed in the cerebellum were also observed in individuals with FHD. We conducted 2 x 2 repeated measures ANOVAs for pre-tapping versus post-tapping BOLD signal, and for pre-tapping versus tapping BOLD signal as described above for CD contrasts, using the same significance thresholds.

#### 3. EMG analyses

Due to the small sample size for EMG data and variability of muscle involvement across CD patients, we did not conduct statistical comparisons between groups, but we evaluated, for visualization purposes, the data in two ways as preliminary, qualitative evidence to drive future larger scale studies.

Group-level EMG data: First, we computed group average data for power spectrum density (PSD) for the SCM channel since signal in this region best mirrored group fMRI time courses in the putamen and cerebellum. Our goal was to show the presence of cervical muscle involvement during the task, rather than to rule out signal contributions from other muscles, such as arm/hand flexors. To extract PSD data we used a frequency range of 20-100 Hz to mitigate motion artifacts and baseline drift (high-pass filter), and to attenuate high-frequency noise (low-pass filter). We averaged PSD over each rest or tapping block for each group and graphed PSD time courses across left-handed tapping runs only, since this was the hand that showed fMRI differences.

Individual-level EMG data: To supplement group analyses, we also examined individual signal plots to seek evidence that sEMG might contribute to BOLD time course features (sustained after movement ceases and escalating as movements are repeated) even at the individual level. For the example CD and control runs shown in the Results, the root mean square was plotted to evaluate sEMG time course characteristics [63] in relation to fMRI BOLD time courses. Specifically, the EMG data were filtered with a 4th order bandpass Butterworth filter with cutoff frequency of 20-200 Hz to mitigate motion artifacts and baseline drift (high-pass filter), and to attenuate high-frequency noise (low-pass filter). This was followed by a 4th order Butterworth notch filter to remove powerline interference at 60 Hz and relative harmonics. Next, the RMS value was computed on the cleaned signal using a non-overlapping window of 5 seconds, across the sequence of the study protocol. This means that each 60-second period of tapping or resting was divided into 12 windows, starting with the beginning of the task. The six transition period (rest-to-tapping-to-rest) TRs were excluded from this analysis, as they were from the fMRI statistical analyses.

## Results

CD participants showed sustained fMRI BOLD signal in both the putamen and cerebellum following repetitive movement of the left (non-dominant) hand, while the original FHD cohort that exhibited putamen signal elevation did not show this effect in the cerebellum. We also observed an increase in BOLD signal at the end, relative to the beginning, of tapping blocks in CD but not in controls. BOLD time courses in primary motor cortex regions corresponding to neck representation showed activation during tapping (relative to rest) in both CD and controls, suggesting there was brain neck representation activity in both groups during tapping. In parallel, EMG amplitude in the right SCM showed persistent escalation over the course of each run that mirrored the “climbing” signal in BOLD time courses. Finally, all group differences in brain activity were observed in the absence of group differences in tapping performance or head movement in the scanner and were not associated with global signal drift.

### 1. Tapping task performance during fMRI acquisition

#### Tapping performance

Button press data recording finger tapping frequency and quantity verified that study participants performed the task properly, i.e., that they tapped during tapping blocks and did not tap during rest blocks. CD and controls did not differ in the number of total button presses per run for either hand (p>0.1), and there was no group (CD versus control) by tapping block number interaction for number of button presses for either left-or right-hand tapping (p>0.1).

#### Hand motion sensor measures during rest blocks

Although button press data confirmed all participants performed the task correctly and did not tap during rest blocks, we also used motion sensors to determine whether any other hand movement took place during rest blocks, specifically to rule out that spurious hand movement might have explained our observed fMRI effects. There were no group (CD versus control) by event (pre-versus post-tapping rest block) interactions for motion sensor signal for either hand in which CD and controls showed fMRI differences (left hand tapping), indicating that CD and controls did not exhibit differences in motion sensor signal amplitude during pre-versus post-tapping rest blocks (p>0.1 for each hand). There was also no signal difference by group alone (p>0.1 for each hand) or event alone (p>0.1 for each hand). Finally, there was no increase in hand movement within either group from pre-to post-tapping rest blocks for either CD or controls (one-tailed, paired samples t-tests, p>0.1).

#### Head movement during imaging

Comparison of the average TR-by-TR head movement vector during rest blocks in CD versus controls indicated that there was no significant difference in head movement across groups during rest blocks for either left-or right-hand tapping (p>0.1; Supplementary Figure 2A). There also was no group (CD versus control) by event (pre-versus post-tapping rest) interaction relating to head movement for either left-or right-hand tapping (p>0.1; Supplementary Figure 2B). Finally, there was no relationship between (1) the difference in head movement between post- and pre-tapping rest blocks and (2) the difference in BOLD signal during post-versus pre-tapping rest blocks in CD participants in the ROIs where we saw our main fMRI effects (left-hand tapping: left putamen: R = -0.178, p = 0.579; left cerebellum: R = 0.157, p = 0.627; Supplementary Figure 2C,D). Controls also did not show a relationship between rest elevation in the right putamen during left hand tapping (R = 0.248, p = 0.438).

### 2. fMRI analyses

#### Pre-tapping rest versus post-tapping rest contrasts

##### CD versus control

###### A priori ROIs

There was a significant group (CD versus controls) by event (pre-versus post-tapping rest) interaction for BOLD signal in the left putamen for left-hand tapping runs, and a trend in the left cerebellum for left-hand tapping runs (Table 2; Figure 1A). There were no CD/control differences in right hemisphere *a priori* ROIs for left-hand tapping or in either hemisphere for right-hand tapping. For the main analysis, there was an effect of condition (pre-tapping rest versus post-tapping rest) independent of group, for the right putamen for left-hand tapping runs (F=17.827, p=0.000350; Figure 1B), but no effects of group or condition for right-hand tapping runs. Post hoc t-tests confirmed that individuals with CD showed significantly greater post-tapping rest block BOLD signal relative to pre-tapping rest block signal in the left putamen and left cerebellum for left hand tapping runs, while controls did not show this difference in either of these regions (Table 2, Figure 1A). The right putamen signal was significantly greater for rest blocks after, versus before, left-hand tapping in CD, and there was also a trend toward significance for this effect in controls (Table 2, Figure 1B). For this same comparison in the right cerebellum, there was a trend toward significance in CD, but no difference in controls (Table 2, Figure 1B).

**Table 2:** Group (CD versus controls) by event (pre-versus post-tapping rest) interaction for BOLD signal for *a priori* ROIs, and posthoc within-group comparisons.

|  | LHem group x<br>event<br>interaction<br>(F/p) | RHem group<br>x event<br>interaction<br>(F/p) | LHem CD<br>Pre vs. post<br>(t/p) | RHem CD<br>Pre vs. post<br>(t/p) |
| --- | --- | --- | --- | --- |
| Left hand tap |  |  |  |  |
| Putamen | **9.164/0.00619\*\*** | 1.180/0.289<sup>†</sup> | **3.24/0.00800\*\*** | **3.23/0.00800\*\*** |
| Cerebellum | 6.367/0.0190* | 1.718/0.203 | **3.78/0.00300\*\*** | 2.639/0.0230* |
| Right hand tap |  |  |  |  |
| Putamen | 0.728/0.403 | 1.0700/0.312 |  |  |
| Cerebellum | 0.426/0.521 | 0.0010/0.979 |  |  |

*a priori* comparisons are shown in the first two columns (2 x 2 repeated measures ANOVAs); the remaining comparisons are posthoc, two-tailed paired samples t-tests to evaluate whether *a priori* findings reflected pre- to post-tapping rest BOLD signal changes in CD. Note that because there was no group or event effect for right-hand tapping runs it was not useful to evaluate events separately in post hoc tests (this finding indicated neither group showed an effect of event). LHem=Left hemisphere; RHem=Right hemisphere; CD=cervical dystonia group; Ctrl=healthy control group; **\*\*significant**; \*trend; <sup>†</sup>showed a significant effect of event (pre-versus post-tapping blocks), but no group x event interaction.

###### Other sensorimotor ROIs

After correcting for multiple comparisons, no other sensorimotor ROIs (including neck representations in primary motor cortex) showed significant (or trend) group by event interactions for pre-versus post-tapping rest BOLD signal.

#### Pre-tapping rest versus tapping contrasts

##### CD versus control

###### A priori ROIs

There was a significant group (CD versus controls) by event (pre-tapping rest versus tapping) interaction for BOLD signal in the left putamen (F=14.428; p=0.00100) and left cerebellum (F=11.453; p=0.00270) for left hand tapping runs (Figure 1A). There was also a trend for the group by event interaction for BOLD signal in the right cerebellum for left-hand tapping runs (F=5.448, p=0.029; Figure 1B). No other conditions/hemispheres in these two *a priori* ROIs showed an interaction.

###### Other sensorimotor ROIs

After correcting for multiple comparisons, no other sensorimotor ROIs (including neck representations in primary motor cortex) showed significant group by event interaction for tapping versus pre-tapping rest BOLD signal for either left- or right-hand tapping. The left area 2 of somatosensory cortex showed a trend during left-hand tapping (F=10.154; p=0.00400).

##### Analysis of signal “escalation” in fMRI time courses during tapping (quantitative testing of qualitative observations made in Blood et al., 2004)

For the two *a priori* ROIs that showed significant group x task differences during tapping in our main contrast analyses (left putamen and left cerebellum) the post hoc “escalation” analysis showed that the BOLD signal in these two ROIs increased over time in CD participants as they repeated the tapping task. Specifically, a two-tailed, paired t-test showed CD exhibited a significant change from the beginning to the end of the block (t=3.422; p=0.0057) for left putamen (Figure 2A), but controls did not (t=1.559; p=0.147). Similarly, for left cerebellum, CD exhibited a change from the beginning to the end of the block (t=3.0847; p=0.0104) but controls did not (t=0.154; p=0.881) (Figure 2B).

**Figure 2.**
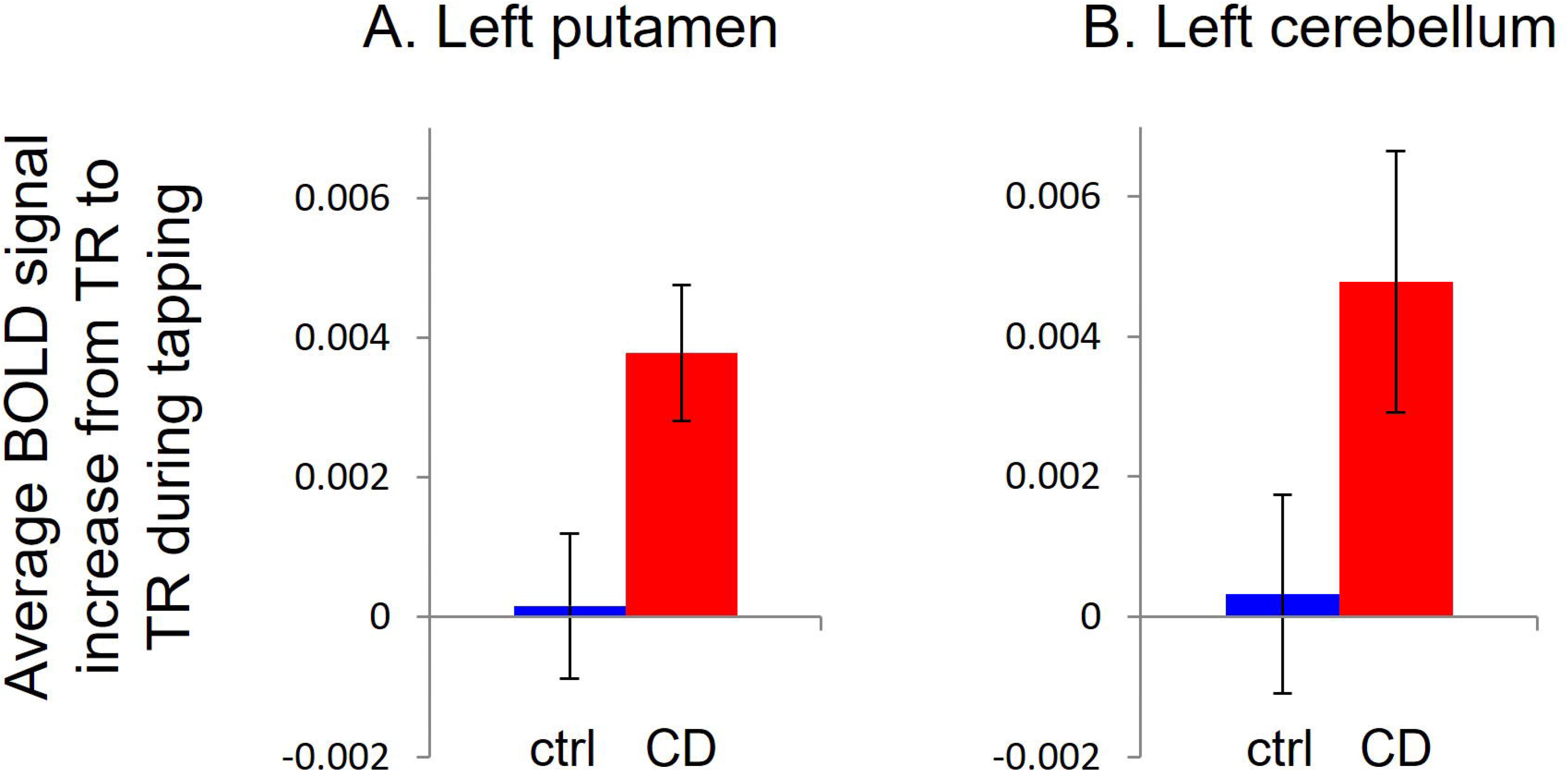
Bar graphs showing average within-subject change in BOLD signal from the beginning (TRs four through seven [first four TRs after steady state signal reached]) to the end (last four TRs) of tapping blocks for CD and controls in (A) left putamen and (B) left cerebellum.

##### fMRI time course analysis in non-activated regions to rule out scanner signal drift (negative control analysis)

Analysis of voxels below activation threshold in ROIs showing significant group differences (left putamen and left cerebellum during left hand tapping) showed no group by event interaction on BOLD signal (p>0.1), and no signal differences between pre-versus post-tapping rest blocks in either control or CD (p>0.1).

##### Post hoc questions: Evaluation of neck representation activation during tapping in both groups (1°Md and 1°Mv)

Both CD and control groups showed BOLD time courses consistent with activation in 1°Md and 1°Mv during tapping relative to rest blocks (Figure 1A, B). Future studies should evaluate this question *a priori* (rather than from already-extracted time course data) to determine if these signals reached statistical significance for activation. In Figure 1A and 1B we show time courses for 1°Md and 1°Mv as a companion to the putamen and cerebellum time course data, and in Figures 3 and 4 we show location of activation during tapping in these regions to demonstrate the consistency of this effect across participants/groups, as well as to show independence of these activation clusters from the hand representation (i.e., nearly all the individual clusters were entirely within the neck representation ROI as shown in Figures 3 and 4). For time course and location images we show the neck region (1°Md versus 1°Mv) for each hemisphere with effects qualitatively most like the putamen.

**Figure 3.**
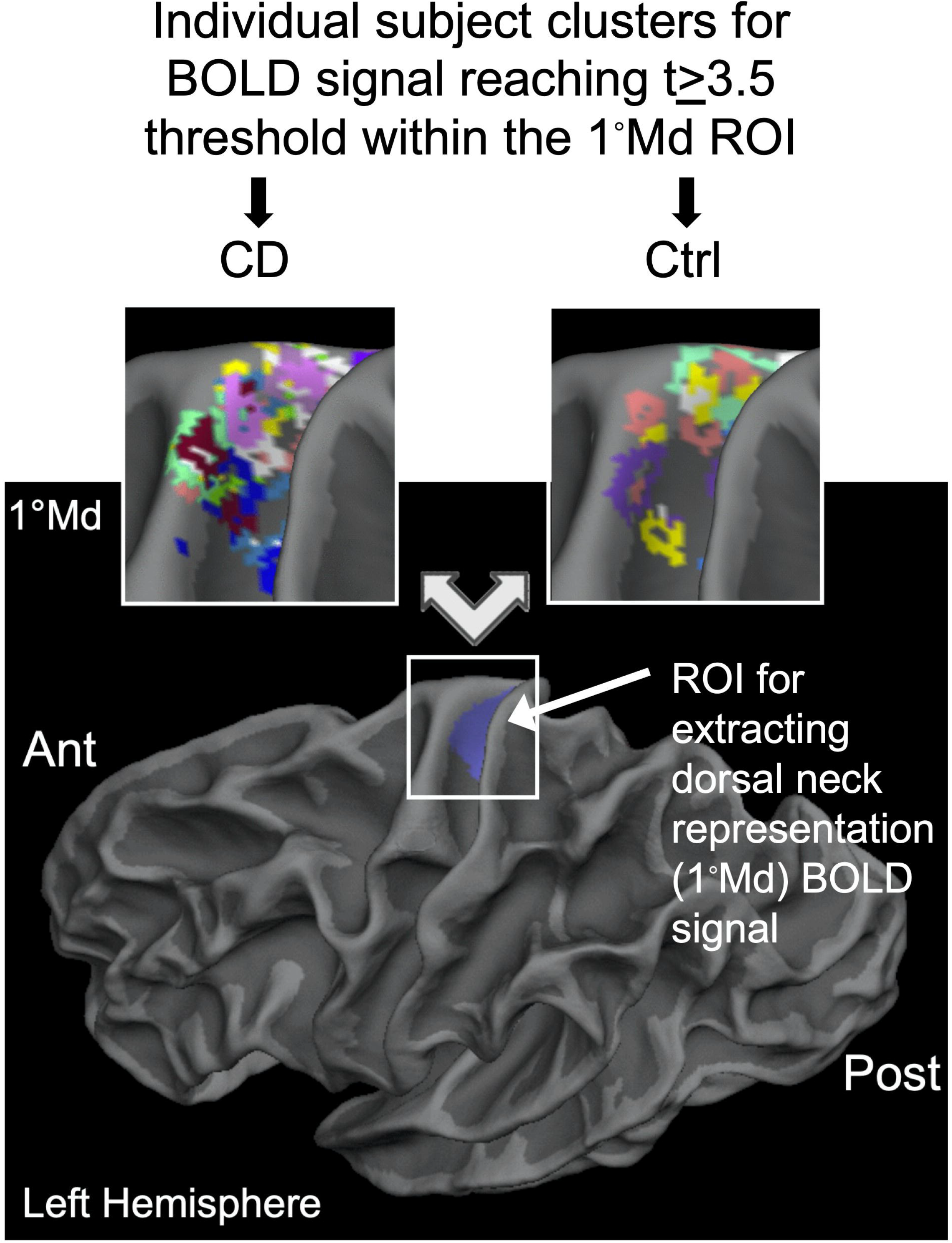
Images showing location of individual cervical dystonia (CD) and control (Ctrl) participant activation in the left dorsal region of primary motor cortex, which encodes representation of the trunk, but has also been shown to include neck representation [60]. Clusters were identified based on the intersection of the 1°Md ROI with t values of 3.5 or greater from the tapping versus rest contrast at the individual (native space) level. These clusters were used in our analyses to extract both rest and tapping-related BOLD signal. The 1°Md ROI is shown in purple on the whole brain, and two enlarged panels showing activation clusters are centered over this region. The hand representation begins immediately ventral to the 1°Md label (see Supplementary Methods for segmentation methods and boundaries). For viewing purposes, outlines of each cluster are shown in a different color for each individual and layered on top of one another. Note that cluster borders in most (but not all) individuals remain within the segmented ROI and thus appear to be distinct clusters in this region (i.e., not “overspill” from the hand region). Clusters are displayed on the white matter contoured surface of the Freesurfer "fsaverage" template brain. Clusters were identified in native space and projected onto the template brain for this figure. Images are viewed using tksurfer software (http://freesurfer.net/fswiki/tksurfer). Ant=anterior; Post=posterior.

In the initial tapping block design fMRI GLM analysis within each participant, 11 of 12 CD, and 7 of 12 controls showed BOLD signal that met our t≥3.5 extraction threshold within the left 1°Md ROI during left hand tapping (Figure 3). 9 of 12 CD, and 10 of 12 controls had BOLD signal met our t≥3.5 extraction threshold in the region just ventral to the hand representation (i.e., in the vicinity of neck representation) within the right 1°Mv ROI during left hand tapping (Figure 4). Of the remaining three CD patients and two controls, two CD and one control showed BOLD signal only in the most ventral part of this ROI (i.e., ventral to neck representation).

**Figure 4.**
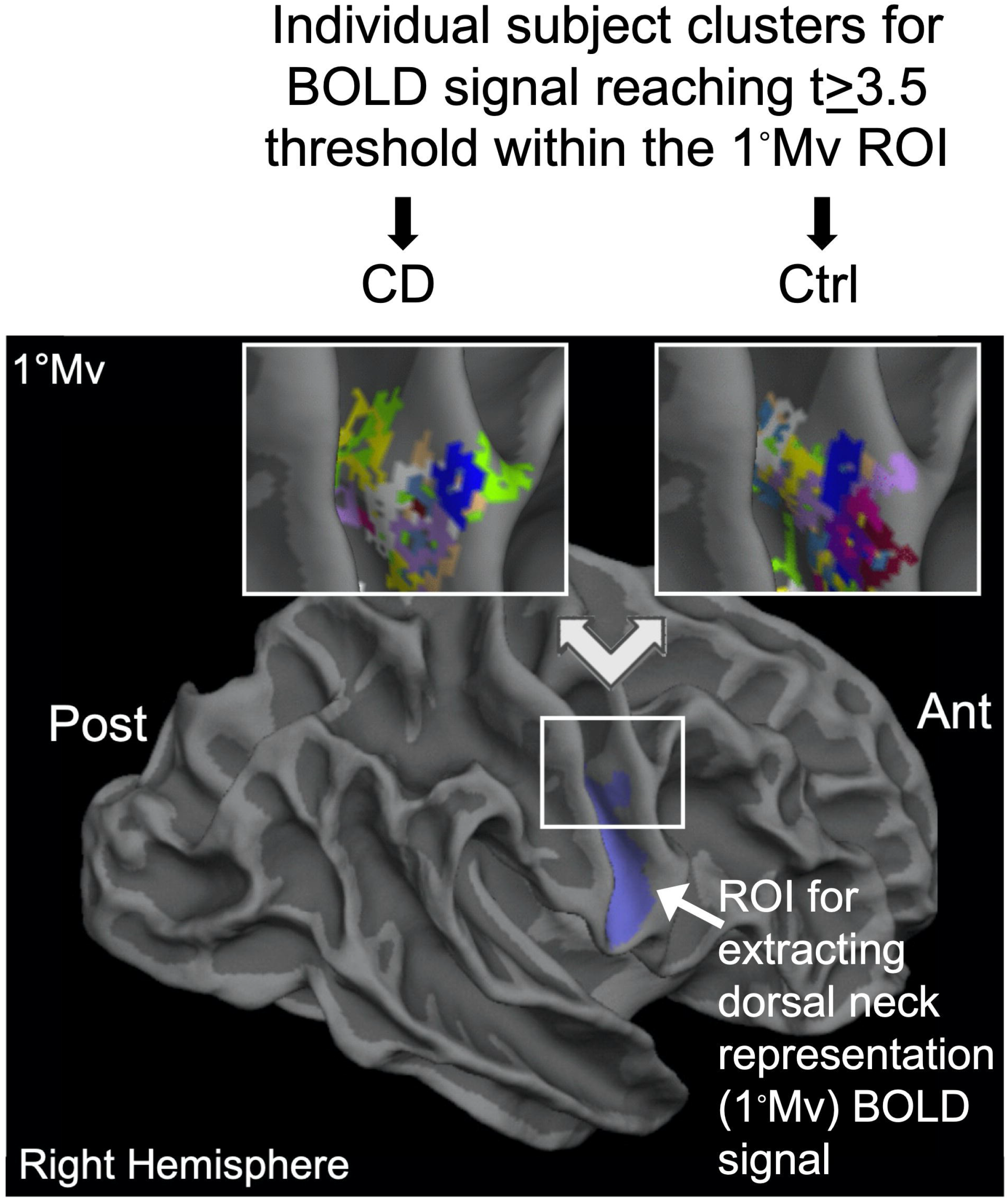
Images showing location of individual cervical dystonia (CD) and control (Ctrl) participant activation in the vicinity of the right primary motor cortex ventral to the hand representation. Clusters were identified based on the intersection of the 1°Mv ROI with t values of 3.5 or greater from the tapping versus rest contrast at the individual (native space) level. These clusters were used in our analyses to extract both rest and tapping-related BOLD signal. The 1°Mv ROI is shown in purple on the whole brain, and two enlarged panels showing activation clusters are centered over the region of primary motor cortex in which neck representation is expected, immediately ventral to the hand/finger representation. The hand representation begins immediately dorsal to the 1°Mv label (see Supplementary Methods for segmentation methods and boundaries). For viewing purposes, outlines of each cluster are shown in a different color for each individual and layered on top of one another. Note that cluster borders in most (but not all) individuals remain within the segmented ROI and thus appear to be distinct clusters in this region (i.e., not “overspill” from the hand region). Clusters are displayed on the white matter contoured surface of the Freesurfer "fsaverage" template brain. Clusters were identified in native space and projected onto the template brain for this figure. Images are viewed using tksurfer software (http://freesurfer.net/fswiki/tksurfer). Ant=anterior; Post=posterior.

##### Post hoc questions: fMRI contrasts from the FHD cohort in our previous publication

*Pre-tapping rest versus post-tapping rest group contrasts for the cerebellum:* There was no significant (or trend-level) group (FHD versus controls) by event (pre-versus post-tapping rest) interaction for BOLD signal in the left or right cerebellum in the FHD cohort from our 2004 publication.

*Pre-tapping rest versus tapping group contrasts for the cerebellum:* There was no significant (or trend-level) group (FHD versus controls) by event (pre-tapping versus tapping) interaction for BOLD signal in the left or right cerebellum for the FHD cohort from our 2004 publication.

### 3. EMG analyses

Group-level EMG data: Group averages indicated that the right SCM muscle most consistently showed abnormal features that mirrored left hemisphere fMRI time courses in CD patients. Group data are shown next to fMRI BOLD time courses in Figures 5 and 6 for comparison. For the left SCM, sEMG time courses were more similar between CD and controls, as were their right hemisphere fMRI BOLD time courses.

**Figure 5.**
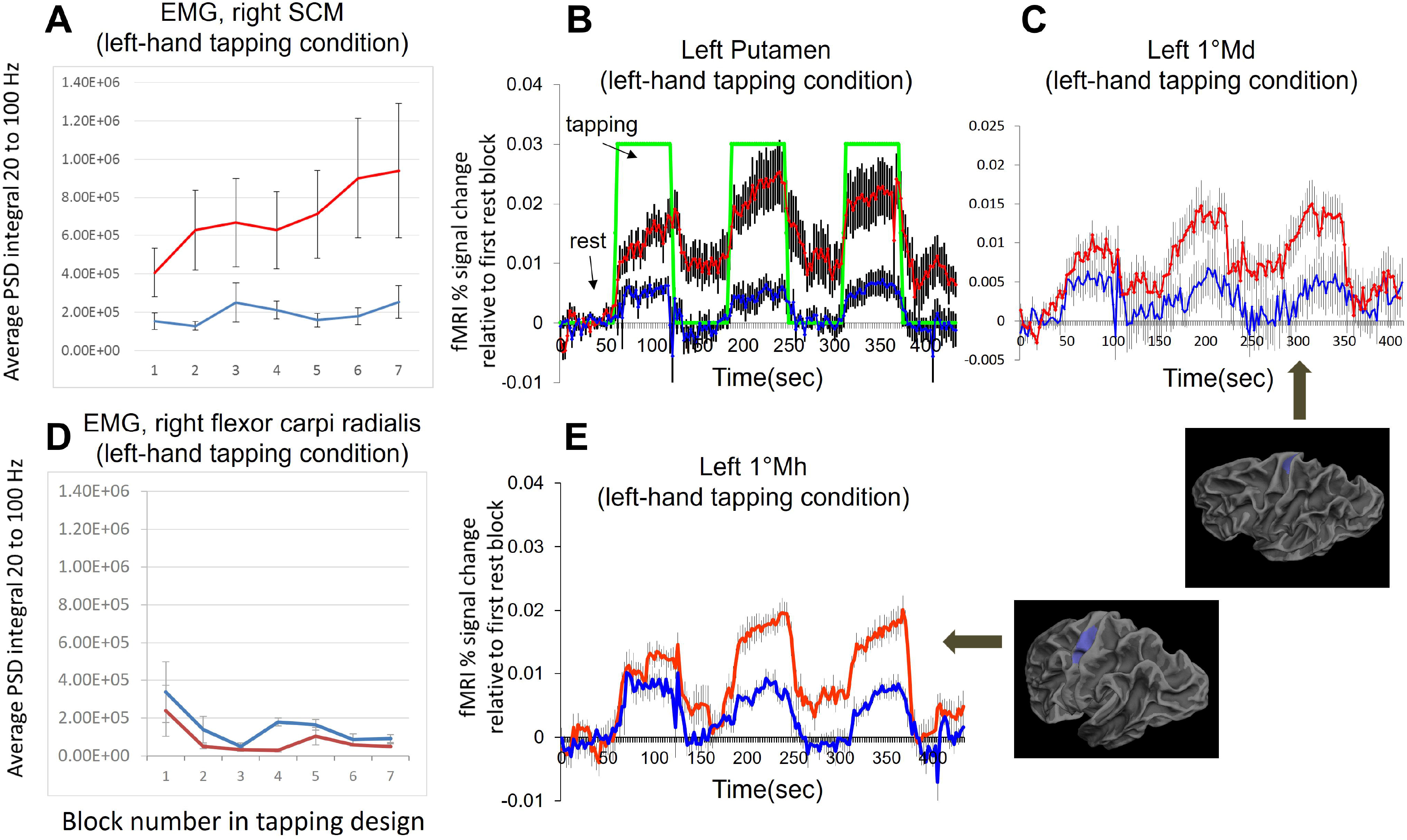
Comparison between right sternocleidomastoid (SCM) muscle EMG and left hemisphere fMRI BOLD signal during left-hand tapping, showing alignment of EMG/fMRI signal patterns. Importantly, fMRI BOLD signal dynamics in the putamen and cerebellum resembled signal traces from cervical muscles. Red corresponds to CD participants and blue to control participants. (A) Right SCM EMG averages for CD and control groups, averaged across each block. PSD integrals were computed by block for the 20 to 100Hz signal frequency range and averaged across the 6 CD and 5 controls with EMG data. (B) fMRI BOLD time courses in left putamen across each run for CD and control groups; data are same graphs as in Fig. 1A. For fMRI BOLD time courses here and also in (C) and (E), data are expressed as percent signal change (decimal equivalent) from the initial, pre-tapping rest block over time. The block design (rest versus tapping) is denoted by the green line on the putamen time course; total run time was 435 seconds, or seven minutes, 15 seconds (with 60 second rest blocks). (C) fMRI BOLD time courses in left dorsal primary motor cortex (1°Md) across each run for CD and control groups. The 3D brain below the BOLD time courses shows the *a priori* 1°Md region-of-interest in blue. For all time courses in this figure, error bars represent standard error of the mean. (B) and (C) data are the same as shown in Fig. 1A. (D) Right flexor carpi radialis averages for CD and control groups, averaged across each block. PSD integrals were computed by block for the frequency range between 20Hz and 100Hz, then averaged across the 6 CD and 5 controls who had EMG data. (E) fMRI BOLD time courses in left hemisphere primary motor cortex (1°Mh) across each run for CD and control groups. The 3D brain to the right of BOLD time courses shows the *a priori* 1°Mh region-of-interest in blue.

**Figure 6.**
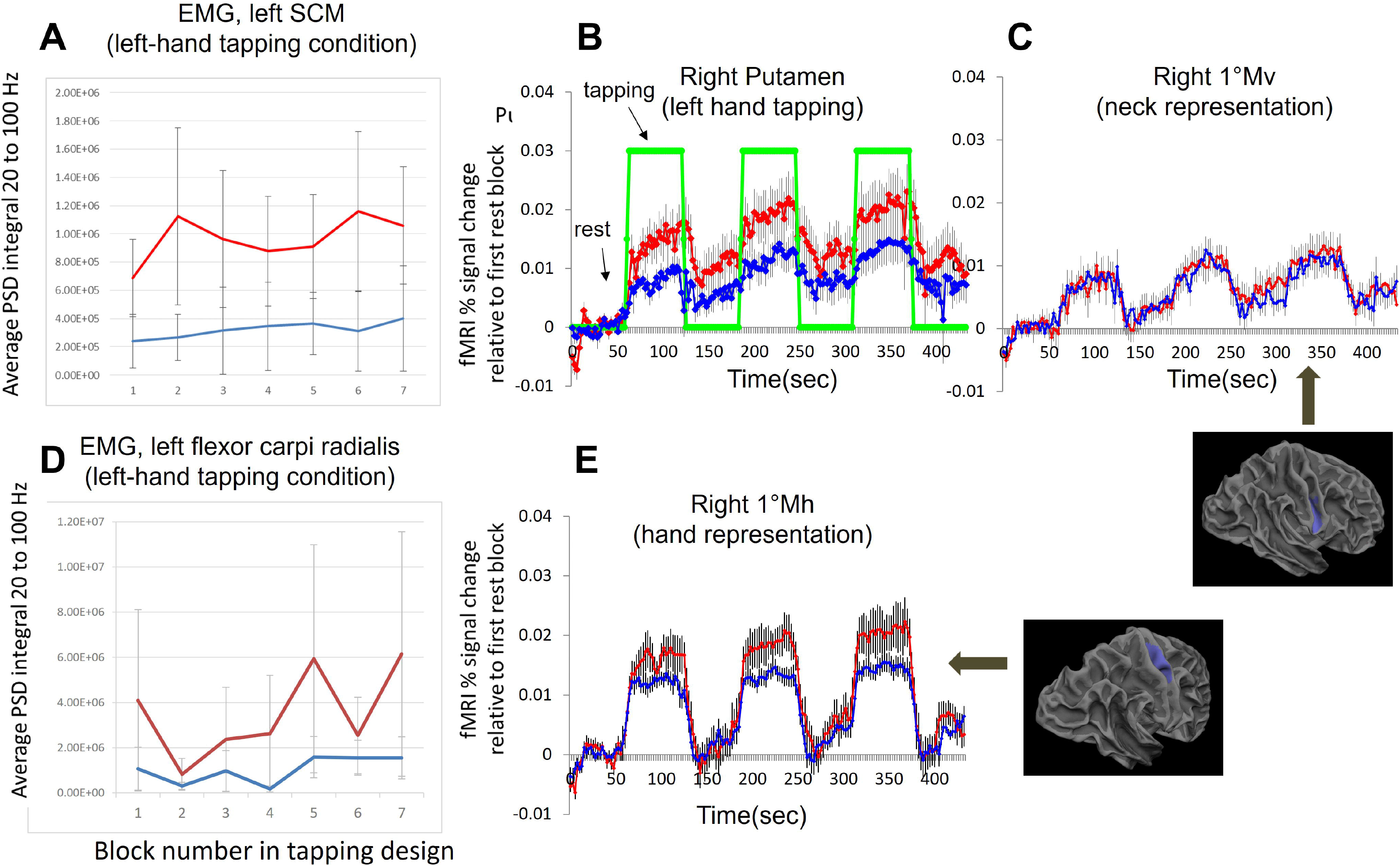
Comparison between left sternocleidomastoid (SCM) muscle EMG and right hemisphere fMRI BOLD signal during left-hand tapping, showing alignment of EMG/fMRI signal patterns. There was greater fMRI BOLD time course similarity between CD and controls in the right than the left hemisphere and this was paralleled by greater between-group similarity of EMG signal traces. Red corresponds to CD participants and blue to control participants. (A) Left SCM EMG averages for CD and control groups, averaged across each block. PSD integrals were computed by block for the 20 to 100Hz signal frequency range and averaged across the 6 CD and 5 controls with EMG data. (B) fMRI BOLD time courses in right putamen across each run for CD and control groups; data are same graphs as shown in Fig. 1A. For fMRI BOLD time courses here and also in (C) and (E), data are expressed as percent signal change (decimal equivalent) from the initial, pre-tapping rest block over time. The block design (rest versus tapping) is denoted by the green line on the putamen time course; total run time was 435 seconds, or seven minutes, 15 seconds (with 60 second rest blocks). (C) fMRI BOLD time courses in right ventral primary motor cortex (1°Mv) across each run for CD and control groups. The 3D brain below the BOLD time courses shows the *a priori* 1°Mv region-of-interest in blue. For all time courses in this figure, error bars represent standard error of the mean. (B) and (C) data are the same as in Fig. 1B. (D) Left flexor carpi radialis EMG averages for CD and control groups, averaged across each block. PSD integrals were computed by block for the frequency range between 20Hz and 100Hz, then averaged across the 6 CD and 5 controls who had EMG data. (E) fMRI BOLD time courses in right hemisphere primary motor cortex (1°Mh) across each run for CD and control groups. The 3D brain to the right of the BOLD time courses shows the *a priori* 1°Mh region of interest in blue.

Individual-level EMG data: Individual sEMG data were also strongly consistent with the imaging data. When compared to controls, patients showed larger amplitude activity in cervical muscles contralateral to the non-tapping hand, as they did also in motor cortex BOLD signals. Features such as (1) escalation of the signal and (2) lack of return to baseline during subsequent rest blocks were observed for cervical sEMG in CD patients just as they were for fMRI time courses, especially in SCM contralateral to the tapping hand (Fig. 7A). In addition, we observed that even though healthy controls showed less of the effects observed in CD participants, they often exhibited some degree of activity change in cervical muscles across tapping task runs (Fig. 7B). There was considerable cross-individual variability in these effects. Such convergence supports the hypothesis that this association was physiologic in nature. Larger sample sizes and tailored study protocols are required to infer causation.

**Figure 7.**
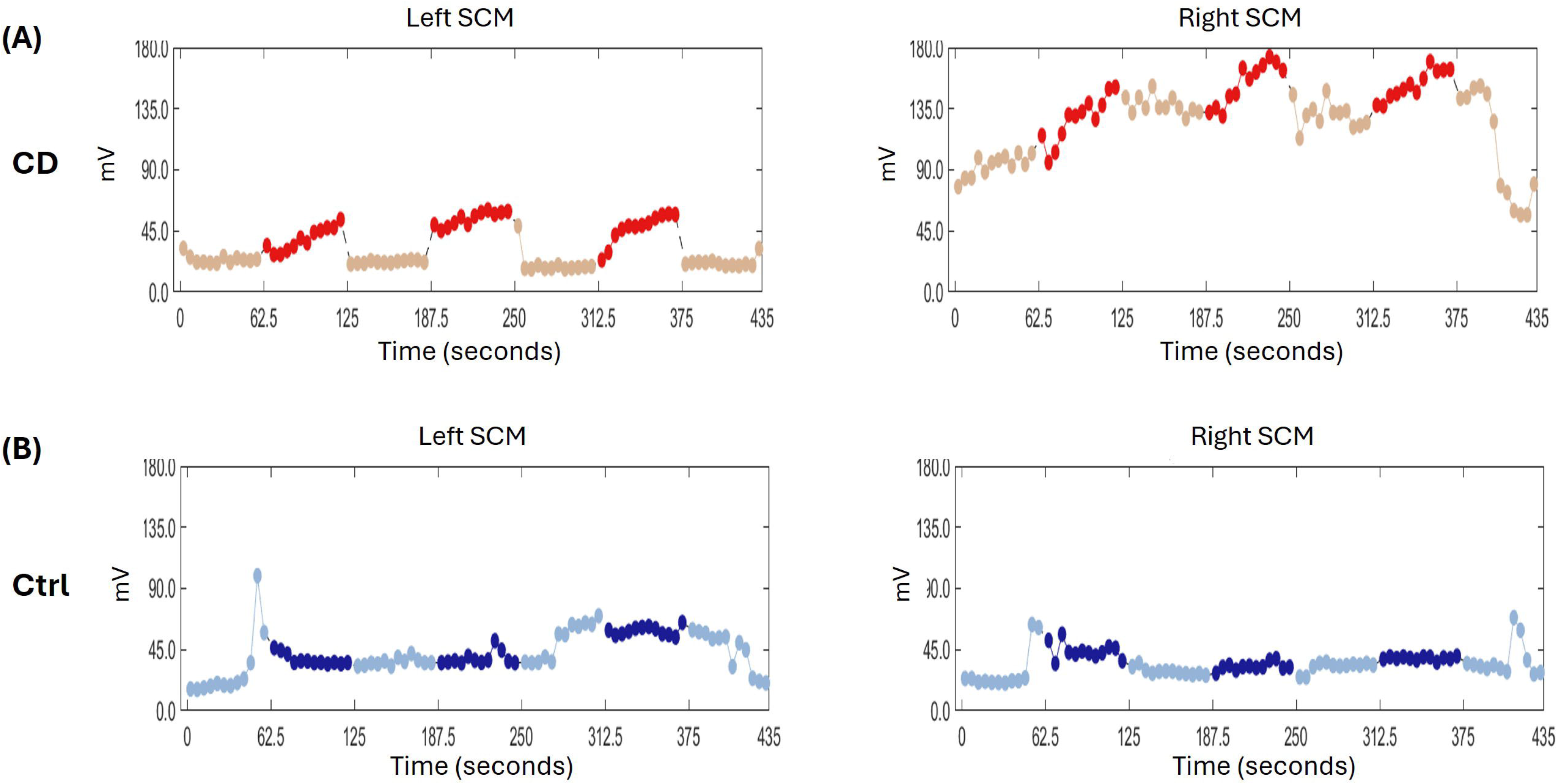
Root mean square (RMS) plots for the sternocleidomastoid (SCM) muscles across a single left-hand tapping run for (A) one CD participant (red/pink) and (B) one control participant (dark blue/light blue). Each marker represents sEMG RMS of the signal amplitude (mV) averaged over a non-overlapping window of 5-seconds. Pink (CD) and light blue (control) denote rest blocks, and red (CD) and dark blue (control) denote tapping blocks.

## Discussion

The findings in this study replicate our previous report of sustained activity in the basal ganglia after repetitive finger tapping in focal hand dystonia (FHD) [10], and extend this observation to CD. This indicates that the effects of muscle/motor program “use”, including repetitive or sustained use of muscles, may lead to pathophysiology in dystonias that are not typically considered task-specific or thought to have a strong use-related pathogenesis. This has significant implications for our understanding of development, expression, and treatment of CD since muscle use (behavior) and function amplitude (physiology) can be modified by interventions such as physical/occupational therapy and botulinum toxin. While previous studies of CD have also shown abnormal activation amplitude in sensorimotor circuitry during motor tasks that do not involve overt movement of the symptomatic muscles, the current study is novel in that it showed (1) that activation abnormalities outlasted movement itself and (2) an increase in activation abnormalities as a task was repeated. Preliminary sEMG data suggest that these abnormalities relate to task-related demands on symptomatic musculature, but this requires confirmation in larger cohorts.

### CD showed both sustained BOLD signal after tapping and an increase in BOLD signal over time during repetitive tapping, relative to controls, in the putamen and cerebellum

One hypothesis about pathophysiology in dystonia is that there may be faulty control of brain inhibitory processes [53; 64; 65; 66; 67; 68; 69]. This has been studied in a variety of ways, including showing impaired intracortical inhibition in focal dystonias with TMS (e.g., [70; 71; 72] and altered spectroscopy measures of GABA function [73; 74; 75]. The finding in the current study that signal escalated over time during tapping itself adds further evidence for this hypothesis in the CD population studied, and is consistent with existing hypotheses about reduced inhibition [53; 64; 65; 66; 67; 68; 69; 70; 71; 72; 73; 74; 75] and/or increased gain [76] of motor circuitry output. While there may have been a relationship between escalating signal during tapping and sustained signal following tapping, this was not tested in the current study, nor did we test whether the sustained signal might be seen after a single movement. Because there are many factors that can impact the balance between excitation and inhibition, it is likely that the specific etiology differs across patients. It would be interesting, in future studies, to combine TMS or spectroscopy measures of GABA function with the fMRI measures in the current study to determine if they are related.

### Sustained activity was observed only during left (non-dominant) hand tapping

Because the tapping task was performed bilaterally in our original study [10] we sought in the current study to determine if effects in dystonia patients were observed for both hands when performing the task unilaterally or only for one hand. We found that the effects were observed only during left-handed (non-dominant hand) tapping. There are several potential interpretations to this finding, which can be interrogated more directly in future studies. Since the non-dominant hand is presumably less dexterous than the dominant hand, the findings may reflect elevated function of mechanisms relating to control and/or effort of movement. Indeed, an interpretation regarding mechanisms of motor control is consistent with our model proposing that the indirect pathway of the basal ganglia coordinates a broadly distributed motor control system that complements movement selection and trajectory systems [34; 35]. As mentioned in the Introduction, this system is proposed to use various forms of mechanical impedance to set the degree of control for a given movement. Such a system would be expected to work harder when the inherent control ability of the anatomy performing the task is lower (e.g., the non-dominant hand) and also when more control is demanded by performance (e.g., the degree of precision demanded by a task, which was not tested here). The findings might also reflect characteristics of motor control laterality in the brain. This latter possibility is discussed in the next paragraph. It should be noted that since all participants here were right-handed it is not clear if the results relate specifically to use of the dominant hand or if they reflect hemispheric differences that are independent of hand dominance. Future studies using left-handed individuals are needed to test this.

### Despite the unilateral nature of the task, sustained putamen function in CD was observed bilaterally

For the hand that showed group differences (left hand), sustained function was observed bilaterally in CD, despite the unilateral nature of the task. Moreover, the main group effects were in the hemisphere ipsilateral to movement, suggesting the function of the ipsilateral hemisphere in movement may be relevant to dystonic pathophysiology. During tapping itself, both patients and controls showed positive BOLD signal in bilateral sensorimotor regions (see Figure 1 for example). This observation is consistent with previous imaging studies; a number of imaging studies in healthy controls report bilateral activation of sensorimotor regions during unilateral motor tasks [e.g., [77; 78; 79; 80; 81]] and across the midline “overflow” as measured by EMG, even in healthy controls, shows strong, individual variation and is affected by factors such as age [82]. Moreover, since sEMG has a low signal-to-noise ratio, it may be that so-called “overflow” plays a role in normal motor control across all individuals; to our knowledge, technology has not advanced sufficiently to test this directly.

Studies finding bilateral brain and sEMG activation during unilateral tasks are consistent with models of motor control such as the Dynamic Dominance Model, which proposes mechanically and functionally distinct specialization of the left versus right hemispheres, each contributing to the control of movement on either side of the body [83; 84]. In addition, Shibuya and colleagues have previously shown that ipsilateral motor cortex function during a unilateral finger motor task was related to the force exerted [78], suggesting that abnormal left putamen elevation observed here in CD may have related to altered or sustained force control of the tapping hand. Because force control has been shown to be altered in dystonia, and specifically to be less variant [85], this suggests future studies could test the hypothesis that the patient/control differences we saw in the left hemisphere would be amplified by increasing the required force of the task. It is also possible that force control itself is accompanied by contralateral or bilateral axial adjustments that counteract the impact of the distal force on proximal stability and that this effect is amplified in dystonia.

*BOLD time courses in primary motor cortex regions that included neck representation and sEMG time courses from cervical muscles suggested task-related motor signals were sent to non-moving, but still working synergistic muscles [14; 16; 17; 26; 27; 28] to stabilize body regions in both CD and controls*.

In contrast with our previous study [10], the effects we observed here were in association with a task that did not involve explicit instructions to move or otherwise involve the anatomy affected by dystonia, namely, the neck. It is possible that our findings reflect a trait of neural or neurovascular coupling physiology differences in dystonia patients that is not directly symptom-related (e.g., [31; 86; 87; 88; 89; 90]). While we do not rule this out in the current study, we raise the additional possibility that the findings reflect implicit demands of the task on the dystonic muscles (i.e., not the moving anatomy, but anatomy potentially involved in background axial muscle function to help stabilize the arm during the hand movements, for example), either with or without symptom expression.

As outlined in the Introduction, the kinesiology-based premise that movement itself is not necessary to recruit symptomatic muscle activity offers a potential explanation for the findings here, including the laterality of the effects we observed and how even controls showed activation in primary motor cortex neck regions during tapping ([14; 16; 17; 26; 27; 28]).. Core stabilizing function during tapping may involve recruitment of bilateral musculature or stability implemented by muscles ipsilateral to movement may trigger a postural reflex in contralateral neck muscles to offset the mechanical effect on the ipsilateral side and would not necessarily lead to changes in body position (i.e., movement), only in body rigidity in resistance to the mechanical perturbation (feedback) and required precision (feedforward) of finger movement. While it is certainly possible that there are hemodynamic response function abnormalities in the CD patients that are independent of brain activity, MRI itself cannot distinguish neurovascular from true task-related brain activity differences and this is precisely why we followed up with EMG.

Both CD and controls appeared to show brain activation outside the hand region during tapping, suggesting that “spread” beyond the moving region of muscle involvement in dystonia may include amplification of a normal phenomenon (e.g., [14; 16; 17; 26; 27; 28; 34; 35]). Also note that the clusters themselves in *a priori* neck representations were not contiguous with the hand representation in most cases for either CD or controls, arguing against the possibility that our findings reflected overspill from the hand region in either group. The sEMG data—including cervical muscle activation in healthy controls during finger tapping—provide a more direct evaluation of our hypothesis than the fMRI data, although the current dataset has limited statistical power due to sample size, along with predicted inter-participant variation. The similarity of qualitative features in sEMG time courses to abnormal fMRI time course features in CD patients warrants consideration in addition to other factors proposed in previous studies.

While sEMG data and analyses evaluating the role of spatially distributed mechanics to brain activation during an anatomically discrete (finger tapping) motor task in the current study were preliminary and exploratory, they offer convergent evidence for a role of task-specific cervical muscle function during finger tapping, and suggest that this additional and/or alternative interpretation should be considered along with and complementary to other interpretations, such as dystonic endophenotypes [29; 30], muscle “overflow” [91], and hemodynamic abnormalities in CD [32]. We emphasize that the EMG observations here are not conclusive; rather, they are hypothesis-generating and point to the potential value of merging mechanical principles from the kinesiology literature with movement disorder models [34; 35].

### The role of the cerebellum in our findings: Do regional differences between findings distinguish between types of dystonia?

There were significant cerebellar differences both during and after tapping in CD that were not observed in the previous FHD cohort. Conversely, in the FHD cohort we observed elevated primary motor cortex BOLD signal during tapping and this was not observed in the CD cohort in the current study. A number of previous studies have shown cerebellar abnormalities in CD [31; 36; 92; 93; 94], including abnormalities that distinguished CD from other forms of dystonia [36]. The findings here thus add to existing evidence that regional differences may be useful in distinguishing different forms of dystonia.

### Methodological considerations of our analyses

As noted in the Introduction, the effects we observed in this study would not have been observed if we had used a typical voxelwise group contrast approach (i.e., without time course extraction and without summing across regions in which patients might show somatotopic reorganization) and when high-pass filtering removes true signal as scanner drift. These findings do not substitute for but, rather, complement studies that deal with localization questions or differences in the topology of activation in dystonia, since the approach we have used does not evaluate this information. However, it should be noted that the findings in the current study should be taken into consideration in localization studies because they have important practical implications for analyzing and interpreting task-related activation. If activity does not return to baseline after a particular population performs a task then this shifts the “control” condition baseline and could potentially lead to false negatives for elevated activation and false positives for reduced activation in a given region.

### Summary

This study identified functional brain abnormalities in CD both during and after repetitive finger tapping that did not involve overt movement of symptomatic muscles, but which may have recruited symptomatic muscle activity as part of background stability control. Specifically, we observed escalating motor function in putamen and cerebellum during the task in CD and sustained function in these regions after completing the task. These effects were observed only during tapping with the non-dominant hand, and group differences were observed only ipsilateral to the tapping hand. The effects in the putamen were similar to sustained function observed previously in FHD patients but differed in that sustained function was also observed in the cerebellum in CD, but not in FHD. sEMG measures from cervical muscles during and after the finger tapping task were consistent with a hypothesis that these findings potentially reflect abnormal elevation of cervical muscle recruitment supporting core mechanical control and stabilization in CD and that repetition of tasks may exacerbate the abnormalities. Future studies using EMG measures in larger cohorts will be needed to determine if and how these abnormalities contribute directly to symptom development and expression. Independent of the source of abnormal BOLD responses, our findings illustrate an important role for muscle “use” and repetitive tasks in potentially reinforcing pathophysiology in at least two distinct forms of dystonia.

