## Supplementary Methods and Supplementary Figures for "Sustained activation in basal ganglia and cerebellum after repetitive movement in cervical dystonia"

1. Normalization procedure for motion sensor data:

Because motion sensor data reflected relative (tapping versus rest), rather than absolute signal, we normalized data values across runs and participants. We wished to normalize by the net amplitude (peak to trough) of the rest signal, so we did this in the following manner: Because values encoded direction of movement associated with moving fingers up and down during tapping, the motion sensor data straddled a zero line and therefore included approximately half negative and half positive values. To calculate the approximate average amplitude from peak-to-trough of “baseline” (rest block signal), we determined the median absolute value across all four rest blocks for each run and hand, and multiplied it by four (absolute value divided the peak-to-trough distance in half, while the median value divided it in half again). We used the median rather than the mean since any remaining large-scale artifacts would disproportionately bias the mean value. We then divided all data points across each run/hand by this “median x four” value.

2. Segmentation of 1°Md and 1°Mv ROI masks for BOLD timecourse extraction

Primary motor cortex for the hand region was defined based on empirical data from [1], identical to our procedures used previously [2]. To investigate other areas of motor cortex that included neck and/or trunk representation ROI masks were defined for extracting data, in regions dorsal [3] and ventral to the hand (1°Mh) region, using the Freesurfer fsaverage template brain and tksurfer viewing program (https://surfer.nmr.mgh.harvard.edu/). The following boundaries were used to define these two masks, keeping in mind when viewing figures of the masks that the deep sulcal ribbon is shown in dark gray, and gyral territory is shown in light gray: The posterior boundary was defined by the center of mass of the central sulcus for M1 regions both ventral (1°Mv) and dorsal (1°Md) to 1°Mh. Because primary motor cortex localizes mostly to sulcal territory in the human (Picard and Strick, 2001), we included only deep sulcal territory in our 1°Md and 1°Mv ROI segmentations to be certain we were as limited as possible to the primary motor region; because our analysis tool extracted any cluster overlapping with this region, this did not limit voxels to the deepest part of the sulcus, it merely constrained the region used for extraction of clusters. The anterior boundary for each of these regions on the inflated brain (which actually represented the most dorsal component of the anterior bank of the central sulcus) was thus defined as the anterior boundary of the sulcal ribbon, detected automatically using tksurfer software (http://freesurfer.net/fswiki/tksurfer). The boundaries abutting 1°Mh were defined by the boundaries of the 1°Mh label itself (detected automatically in tksurfer software). The dorsal boundary for 1°Md was defined at the most dorsomedial region of the central sulcus, just before turning in toward the medial surface of the hemisphere. The ventral boundary for 1°Mv was at the most ventrolateral region of frontal cortex. Images of ROI masks are shown in Supplementary Figure 1 and in main text Figures 3 and 4. Note that because we did not include a localizer neck task to define neck regions, we included territory of other representations in the 1°Md and 1°Mv masks; the primary goal was to extract any activation in motor cortex that was outside representation of the overtly moving anatomy (the hand). However, figures depicting individual localization of activation clusters indicated activity was localized to the region most likely to correspond to neck representation (the dorsal half of the 1°Mv ROI, for example).

3. Head movement during imaging

In addition to standard procedures for head stabilization during data acquisition, and motion correction during fMRI analyses, we used two other layers of quality assurance to be certain head movement was not a factor in our findings (QA Analysis 3, Table 2). First, we screened for and eliminated timecourses with large spikes consistent with residual head movement artifacts remaining after motion correction; procedures and criteria for this step are described below in *Quality assurance preceding group analyses of fMRI timecourse data*.

Second, for task conditions showing group differences, we used motion correction values from our image preprocessing step (see below) to evaluate whether there were (1) group (CD versus control) differences in head movement during rest blocks, which were the main event(s) of interest, and if there were (2) group by event (pre- versus post-tapping rest blocks) interactions of head movement. For the first test we used a two-tailed t test, and for the second we used a 2 x 2 repeated measures ANOVA, with event (pre- versus post-tapping rest) as the repeated measures factor. This evaluated head movement data for potential group x event interactions, analogous to the comparisons made for BOLD signal (see below). Data was extracted for these analyses in the following manner: We used the total motion vector value output from motion correction parameters as our measure, as it best represented the net amount of movement per participant. For each run, we used these numbers to calculate the amount of movement made from one TR to the next (inter-TR difference), and used absolute values of these differences to compute an average amount of movement across all TRs for each event of interest. Average values per run were then averaged across runs for each participant/condition before numbers were entered into each analysis.

In addition to evaluating whether there were group differences in head movement, we ran a negative control analysis testing more directly whether pre- versus post-tapping rest block BOLD signal differences in CD participants could be explained by differences in head movement during pre- versus post-tapping blocks. Specifically, we calculated pre- versus post-tapping head movement differences for each participant (using the average values described in the paragraph above), and evaluated the relationship of these values to pre- versus post-tapping BOLD signal differences, using a linear regression.

**Supplementary Figure Legends**

Supplementary Figure 1. Images showing ROI segmentations on the inflated Freesurfer "fsaverage" template brain for the left (A) and right (B) hemispheres for primary motor cortex dorsal to the hand representation (1°Md) and primary motor cortex ventral to the hand representation (1°Mv). ROIs are shown in purple. Ant=anterior; Post=posterior. ROIs were used to identify and extract BOLD signal timecourses across each run above threshold in these regions. Images are viewed using tksurfer software (http://freesurfer.net/fswiki/tksurfer).

Supplementary Figure 2. Head movement comparison between CD (red) and controls (blue), and absence of relationship to post-tapping rest block elevation. (A) shows a CD/control comparison of the average inter-TR vector of head movement for all four rest blocks; there were no statistical differences across groups. (B) shows a comparison of the difference (subtraction) between (i) the average inter-TR vector of head movement for post-tapping rest blocks and (ii) the average inter-TR vector of head movement for the pre-tapping rest block. This comparison also showed no interaction between group and event, nor did it show effects for group alone or event alone. This result indicated that elevated post-tapping function observed in CD participants could not be attributed to increased head movement during post-tapping blocks, relative to controls. (C) Scatterplots of individual CD data (n=12), showing the relationship between the post- versus pre-tapping head movement difference [as described in (B)] and the post- versus pre-tapping BOLD signal difference in the left putamen for the left hand tapping condition. (D) Scatterplots of individual CD data (n=12), showing the relationship between the post- versus pre-tapping head movement difference [as described in (B)] and the post- versus pre-tapping BOLD signal difference in the left cerebellum for the left hand tapping condition.

Supplementary Figure 3. Surface EMG (sEMG) was measured bilaterally for all regions of interest. Electrodes were placed over the following muscles: (A) Flexor carpi radialis, (B) sternocleidomastoid (SCM), (C) splenius capitus, and region of overlap for trapezius (B) and (C) the underlying levator muscle. Red dots on the image illustrate the placement of the active electrode. The second (differential) electrode was always placed one inch below (or proximal to, in the case of flexors) the active electrode.
