## Supplementary figures and images for "Sustained activation in basal ganglia and cerebellum after repetitive movement in cervical dystonia"

### Supplementary Figure 1

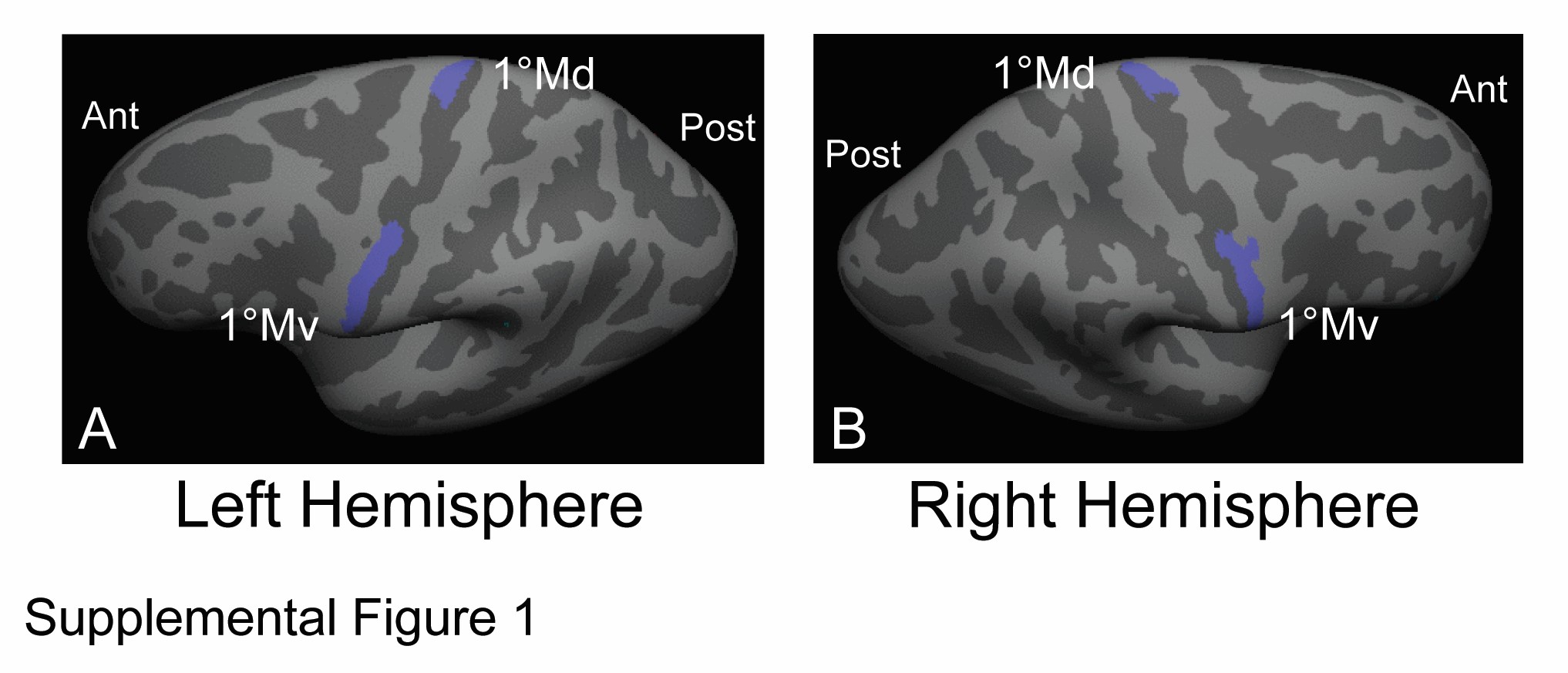

### Supplementary Figure 2

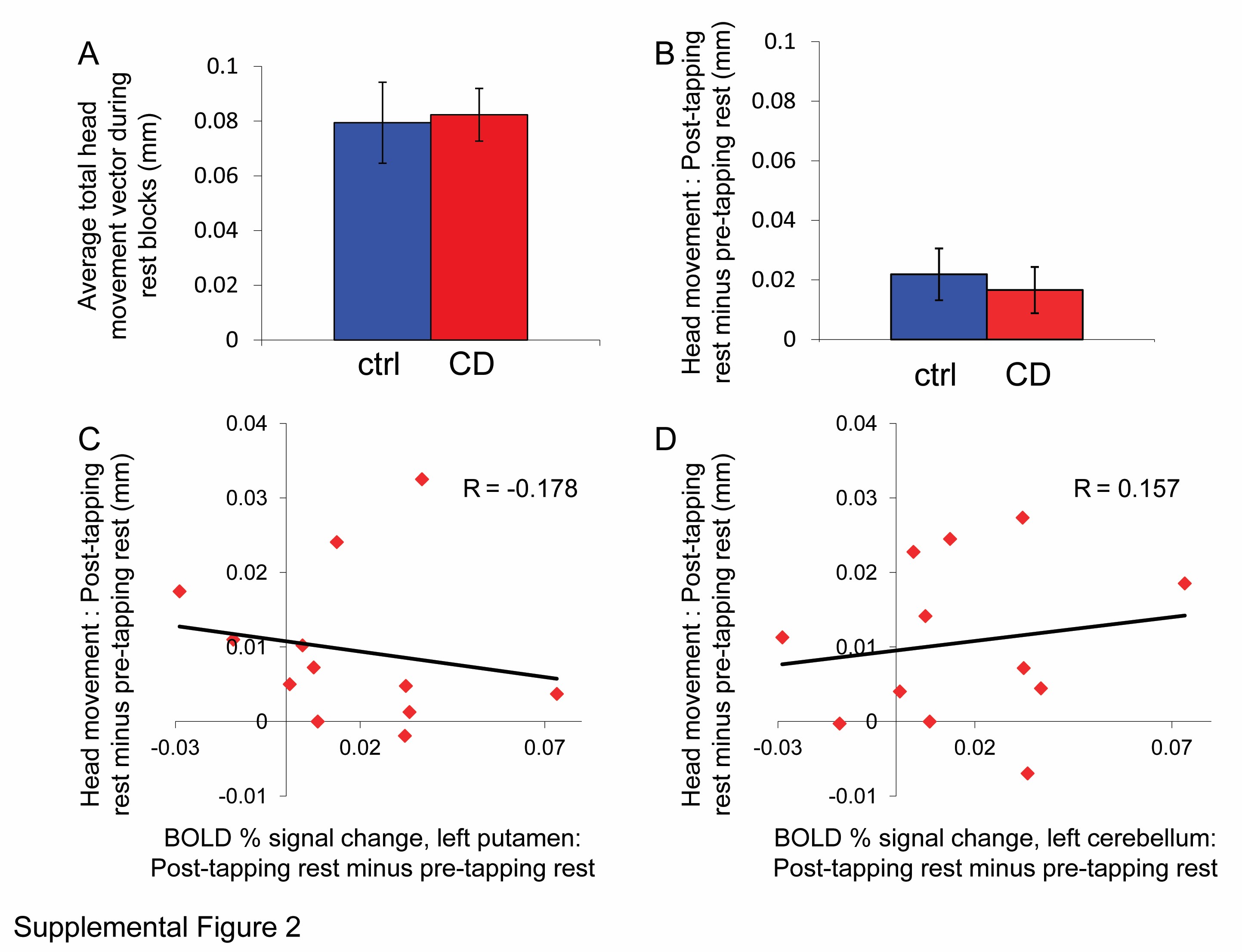

### Supplementary Figure 3

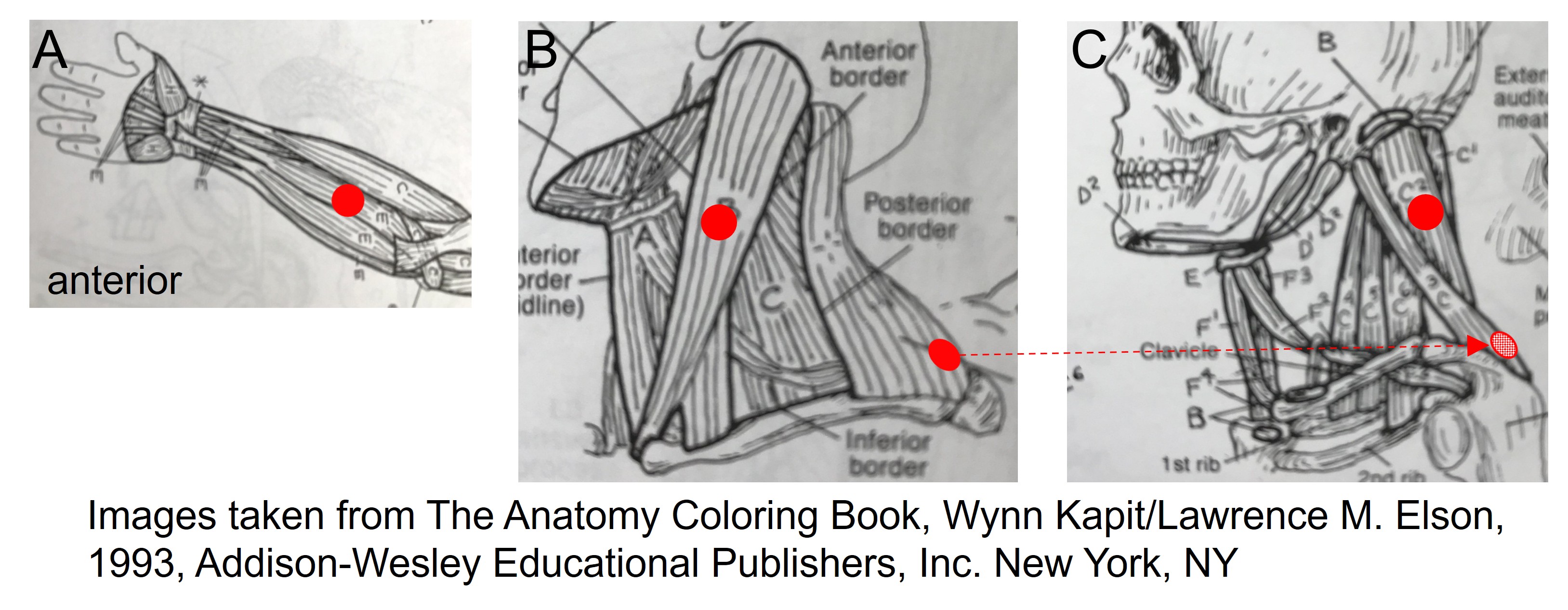
